# Selective Translation Complex Profiling Reveals Staged Initiation and Co-translational Assembly of Initiation Factor Complexes

**DOI:** 10.1101/806125

**Authors:** Susan Wagner, Anna Herrmannová, Vladislava Hronová, Neelam Sen, Ross D. Hannan, Alan G. Hinnebusch, Nikolay E. Shirokikh, Thomas Preiss, Leoš Shivaya Valášek

**Author notes:** Senior Author (Ross D. Hannan, Alan G. Hinnebusch).

## Abstract

Translational control targeting mainly the initiation phase is central to the regulation of gene expression. Understanding all of its aspects requires substantial technological advancements. Here we modified yeast Translational Complex Profile sequencing (TCP-seq), related to ribosome profiling, and adopted it for mammalian cells. Human TCP-seq, capable of capturing footprints of 40S subunits (40Ses) in addition to 80S ribosomes (80Ses), revealed that mammalian and yeast 40Ses distribute similarly across 5’UTRs indicating considerable evolutionary conservation. We further developed a variation called Selective TCP-seq (Sel-TCP-seq) enabling selection for 40Ses and 80Ses associated with an immuno-targeted factor in yeast and human. Sel-TCP-seq demonstrated that eIF2 and eIF3 travel along 5’UTRs with scanning 40Ses to successively dissociate upon start codon recognition. Manifesting the Sel-TCP-seq versatility for gene expression studies, we also identified four initiating 48S conformational intermediates, provided novel insights into *ATF4* and *GCN4* mRNA translational control, and demonstrated co-translational assembly of initiation factor complexes.

## INTRODUCTION

Changing environmental conditions require rapid modifications of gene expression to enable cells to adapt and survive. Regulation of messenger RNA (mRNA) translation – translational control – is central to this complex response (Sonenberg and Hinnebusch, 2009). The initiation phase of translation is the main target of various regulatory inputs that promote or attenuate some of its steps to allow for reprogramming of gene expression. Therefore, it is of fundamental importance to fully understand the molecular mechanism of all initiation steps in the context of their regulatory potential.

The ultimate purpose of the initiation phase is to bring mRNA to the small ribosomal subunit (40S) in such a way that the start of its coding sequence (CDS) is properly identified by the initiator methionyl tRNA (Met-tRNA_i_^Met^) (reviewed in (Hinnebusch, 2014; Shirokikh and Preiss, 2018; Valášek, 2012)). In eukaryotes, the Met-tRNA_i_^Met^ is delivered to the ribosome by the translation initiation factor 2 (eIF2) bound to a GTP molecule in the form of a ternary complex (eIF2-TC). This step is further promoted by eIFs 1, 1A, 3 and 5, producing the 43S pre-initiation complex (PIC). In fact, eIF2-TC and eIFs 1, 3 and 5 can assemble in a higher order protein complex called the multifactor complex (MFC), ensuring more efficient delivery of Met-tRNA_i_^Met^ to the 40S (Asano et al., 2001a; Dennis et al., 2009; Sokabe et al., 2011). All eIFs mentioned thus far also co-operate to prepare the 40S for mRNA docking and subsequent scanning of the 5’ untranslated region (UTR) (Hinnebusch, 2017). The eIF4F cap-binding complex, in cooperation with eIF3, loads mRNA to the 43S PIC to form the 48S PIC. In a scanning-conducive, open conformation, with the anticodon of Met-tRNA_i_^Met^ in the ribosomal P-site in a P^OUT^ conformation, the 48S PIC inspects the mRNA 5’UTR for the proper initiation site. Upon start codon (typically AUG) recognition, accompanied by GTP hydrolysis on eIF2, eIFs 1, 1A, 2, 3, and 5 promote a series of rearrangements leading to a scanning-arrested conformation of the 48S PIC. This entails complete accommodation of Met-tRNA_i_^Met^ in the ribosomal P-site (P^IN^ conformation), closure of the 40S mRNA binding channel and ejection of the eIFs, with the exception of eIF1A and eIF3. While most eIFs are well understood biochemically, information is sparse about when and where along the mRNA 5’UTR interactions among eIFs, mRNA and ribosomal subunits are established and broken during the steps of initiation.

The eIF3 complex, composed of five subunits in yeast (a/Tif32, b/Prt1, c/Nip1, g/Tif35, i/Tif34) and twelve in mammals (a, b, c, d, e, f, g, h, i, k, l, m) (Figure S1A,B), is known to be critical for efficient progression of most of the initiation steps, yet its complete structure has not been determined from any organism (reviewed in (Cate, 2017; Valasek et al., 2017)). The assembly pathway for human and *N. crassa* (similar in composition to human) eIF3 was recently described (Smith et al., 2016; Wagner et al., 2014; Wagner et al., 2016) but it remains to be examined for the most extensively studied *S. cerevisiae* eIF3 complex (Zeman et al., 2019). Recent structural studies of various PICs revealed several well-resolved but otherwise discontinuous densities attributed to various eIF3 modules that together nearly embrace the entire 40S. The yeast a/Tif32-c/Nip1 module, as well as the mammalian eIF3 octamer (a-c-e-f-h-k-l-m), were found to stably reside near the mRNA exit channel, whereas the b/Prt1-i/Tif34-g/Tif35 subunits, as well as their mammalian orthologs (b-i-g), were shown to comprise a module in contact with the mRNA entry channel, where they can interact with incoming mRNA to influence scanning (Figure S1C) (Aitken et al., 2016; Cuchalová et al., 2010; des Georges et al., 2015; Erzberger et al., 2014; Chiu et al., 2010; Llacer et al., 2015). Both modules are flexibly connected by the poorly resolved extended C-terminal half of the eIF3a subunit, allowing the quaternary module (b-i-g-a–Cter.) to re-locate to the interface surface of the 40S subunit, and back, at stages of initiation yet to be fully identified (Figure S1D) (Llacer et al., 2018; Simonetti et al., 2016; Valasek et al., 2017). Importantly, besides initiation, eIF3 was also shown to i) control translation termination and ribosomal recycling (Beznosková et al., 2013; Beznoskova et al., 2015; Pisarev et al., 2007); and to ii) stimulate reinitiation (REI) on downstream cistrons after translation of short upstream uORFs – due to its ability to remain bound to elongating ribosomes immediately following termination (Mohammad et al., 2017; Park et al., 2001; Szamecz et al., 2008). Owing to the manifold functions of eIF3, deregulated eIF3 expression is associated with numerous pathological conditions (reviewed in (Gomes-Duarte et al., 2018; Robichaud and Sonenberg, 2017; Valasek et al., 2017)).

Translation REI can occur on mRNAs carrying more than one ORF, involving either post-termination 40Ses or 80Ses, and is highly regulated in response to various stresses and other intra-or extracellular signals (Gunisova et al., 2018). The mammalian master transcriptional activator *ATF4* and its yeast functional orthologue *GCN4* are among the most studied examples of genes whose expression is regulated by REI in response to stress. The *GCN4* mRNA 5’UTR contains four very short AUG-initiated uORFs preceded by one longer uORF that can be initiated by either of two consecutive non-AUG codons in the same frame (creating nAuORF1 and 2) (Gunisova et al., 2018) (Figure S2A). Both nAuORF1 and 2 were reported to be ribosome-occupied (Ingolia et al., 2009). Nevertheless, translation of only nAuORF2 was independently confirmed, and still appeared to exert no influence on *GCN4* translational control (Zhang and Hinnebusch, 2011). Instead, *GCN4* expression is regulated by the so-called delayed REI mechanism, enacted by the AUG-initiated uORFs 1 through 4, that is exquisitely sensitive to eIF2-TC availability (Gunisova and Valasek, 2014; Hinnebusch, 2005). The 5’ proximal uORF1 and uORF2 are positive, REI-promoting sequences, whose ability to allow efficient REI is determined by five *cis*-acting REI-promoting elements (RPEs i. to v.) mapping upstream of these uORFs and making contacts with eIF3 (Figure S2A). By contrast, the 5’ distal uORF3 and uORF4 are negative, REI-non-permissive sequences lacking any RPEs (Gunisova et al., 2016; Munzarová et al., 2011).

In analogy to *GCN4’s* uORFs, reporter expression experiments with the mouse *ATF4* mRNA 5’UTR revealed that uORF1 is a positive, stimulatory sequence allowing efficient REI after its translation, whereas translation of uORF2 inhibits *ATF4* expression (Vattem and Wek, 2004). Translation of uORF1 combined with low levels of the eIF2-TC is then required to overcome the uORF2 inhibitory effect. Similarly to *GCN4’s* uORF1 and uORF2, the *ATF4’s* uORF1 is surrounded by *cis*-acting, REI-promoting sequences. The upstream sequences most probably form a specific secondary structure contacting eIF3 (Hronova et al., 2017) (Figure S2B). In contrast to mouse *ATF4*, human *ATF4* also possesses an additional AUG-initiated uORF upstream of uORF1 referred to as uORF0 (Lu et al., 2004)), whose role has not been examined. Extensive genetic and biochemical efforts by numerous labs have been devoted to fully grasp the molecular details of these canonical examples of gene-specific translational control; however, further progress has been hindered by the limitations of standard *ex vivo* approaches.

A breakthrough in this direction has been the development of ribosome profiling (Ribo-seq) to study protein synthesis in living cells. Cell lysates are typically prepared in the presence of the elongation inhibitor cycloheximide to preserve polysomes, treated with RNase I, and regions of protected mRNA, i.e. the ribosome footprints (FPs), are isolated bound to ribosomes and identified by high-throughput sequencing (Ingolia et al., 2009; McGlincy and Ingolia, 2017). Ribo-seq gives access to many aspects of translation and even some coupled events such as protein folding and localization (reviewed in (Ingolia et al., 2019)). Translation Complex Profile sequencing (TCP-seq) extends the Ribo-seq approach in yet another direction. Here, rapid formaldehyde fixation of live yeast cells is used to crosslink any type of ribosomal complex bound to mRNA at native positions. The separate purification and sequencing of footprints from both 40Ses and 80Ses adds an ability to survey all mRNA-associated steps of translation, including, importantly and uniquely, the initiation phase (Archer et al., 2016).

In this study, we developed a TCP-seq protocol for human cells, using HEK293T human embryonic kidney cells as a model cell line to characterize human 40S and 80S FPs. We further devised selective (Sel-)TCP-seq for both yeast and human cells, by adding a prior step of immunopurifying ribosomes associated with an initiation factor of interest (FOI). We then compared FOI-associated 40S FPs (FOI::40S FPs) to those of unselected 40Ses to study the association of eIF3 and eIF2 with 40Ses globally, and on selected mRNA 5’UTRs such as *GCN4* and *ATF4*. Finally, we also analyzed FOI::80S FPs to assess co-translational assembly of factors within the yeast MFC.

## RESULTS AND DISCUSSION

### Translation Complex Profile sequencing (TCP-seq) for yeast and human cells

The methodology development for this study commenced independently of the original TCP-seq work, but nevertheless yielded largely convergent approaches for the yeast protocol (for details see STAR Methods and compare with (Shirokikh et al., 2017)). A notable difference is that the published version included an initial separation of mono-and poly-ribosomal complexes from mRNPs and ribosomal subunits not associated with mRNAs, which was omitted here (see Figure 1A). For human TCP-seq, HEK293T cell cultures were incubated with formaldehyde at 0.3% (w/v) for 5 min, followed by detergent lysis, digestion with RNase I for 30 min to release 40Ses and 80Ses from polysomes, and sucrose density gradient ultracentrifugation to separate them. These conditions were chosen to effectively crosslink polysomes while avoiding failure of cell lysis (Figure S3A), and to achieve subsequent conversion of most polysomes into well-separated 40S and 80S fractions (Figure S3B). Separate 40S and 80S FP cDNA libraries were then generated, allowing for a wide range of insert sizes, and sequenced in parallel with standard RNA-seq analysis of total RNA from the same cell lysates (see Table 1 for an overview of generated libraries). Note that formaldehyde crosslinking unavoidably introduced some variability, especially with HEK293T cells, where even small changes in formaldehyde concentration noticeably affected the polysome profile (Figure S3A). As discussed below, this means that some differences in recorded FP sizes, quantity and distribution, especially when comparing between cell types, might have partly technical origins. With that caveat, sequencing and mapping of the resultant libraries consistently gave 40S and 80S FPs with the expected characteristics (see below) and with FP coverage per mRNA showing good correlation between biological replicates for both human (Figure S3C,D) and yeast samples (Figure S3E-J).

**Figure 1.**
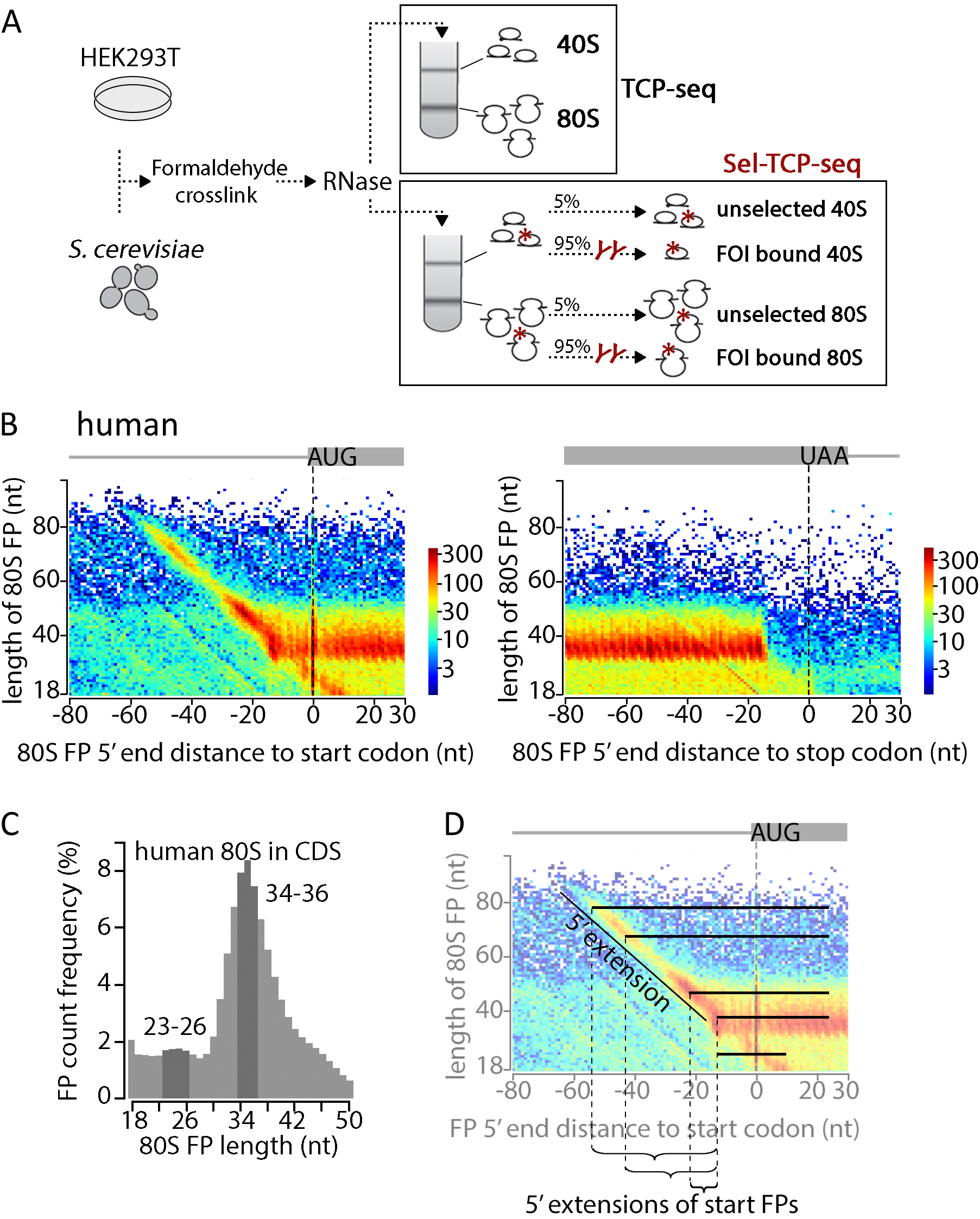
TCP-seq in human cells – full ribosome footprints. (A) Outline of Translation Complex Profile sequencing (TCP-seq) and Selective (Sel)-TCP-seq approaches. 40S, small ribosomal subunit. 80S, ribosome. FOI, factor of interest. See STAR Methods for details. (B) Metagene plot of human 80S footprint (FP) 5’ ends *versus* FP length. 80S FP 5’ ends are given relative to the first nucleotide (position 0) of start (left) or stop codons (right). Colour bars to the right of each plot indicate FP count in log10 scale. Data from 10,778 start sites and 16,803 stop sites are shown. (C) Length distribution of human 80S FPs assigned to annotated coding sequences (CDS; excluding start and stop codon associated FPs) for 9,241 genes is shown. (D) Schematic to facilitate interpretation of plots in (B). Bars display the actual FP sizes that are represented in the heatmap by their 5’ end only. See also Figure S5A,B.

**Table 1.** Libraries overview.

|  | repli-<br>cate | TCP-seq 80S | TCP-seq 40S | Sel-TCP-seq 40S |  |  |  | 80S |  |
| --- | --- | --- | --- | --- | --- | --- | --- | --- | --- |
| | | | | eIF3b | eIF3a | eIF3c | eIF2β | eIF3a | eIF3c |
| Human<br>HEK293T | 1 | x | x | x |  |  |  |  |  |
|  | 2 | x | x |  |  |  |  |  |  |
| Yeast<br>S.c. | 1-1 | x | x |  | x |  |  | x |  |
|  | 1-2 | x | x |  |  | x |  |  | x |
|  | 1-3 | x | x |  |  |  | x |  |  |
| smaller | 2 | x |  |  |  |  |  | x |  |
| smaller | 3 | x |  |  |  |  |  | x |  |

Focusing first on human 80S data, metagene plots show that 80S FPs primarily map within the coding sequence (CDS) of mRNAs and exhibit three-nucleotide periodicity in position (Figure 1B, see Figure S4A for replicate data), as would be expected for translating 80S ribosomes. The predominant human 80S FP type detected by TCP-seq is slightly longer (depending on the replicate, between ∼34-36 nt, Figure 1C or ∼31-33 nt, Figure S4E) compared to the ∼28/29 nt FP length typically seen with conventional Ribo-seq (McGlincy and Ingolia, 2017). This is also somewhat larger than the 80S FP size measured by TCP-seq in yeast (∼29/30nt measured here, Figure S4B-D,F-H and ∼30/31 nt in (Archer et al., 2016)). The small differences in FP size between studies are likely due to variable efficiency of formaldehyde cross-linking and its impact on RNase I accessibility. The same effect likely also accounts for the variability in FP size between replicates. TCP-seq further detects a less abundant, smaller 80S FP type, both in human and more distinctly in yeast (∼23-26 nt and 21/-22 nt, respectively; Figure 1B,C and Figure S4). This smaller 80S FP is thought to originate from 80Ses with an empty A-site and was also observed in ribosomal profiling studies, for example when omitting the cycloheximide pre-treatment (Lareau et al., 2014; Wu et al., 2019). Of note, a proportion of 80Ses residing at the start codon produce substantially 5’ extended FPs, protecting a stretch of up to ∼35 nt emerging from the exit channel (Figure 1B,D and Figure S4A). This is observable to a lesser degree also in yeast TCP-seq (Figure S4B-D). It can be explained by residual binding of eIFs at the exit channel or by ‘queuing’ of a 40S, which were not cleaved off during RNase I treatment due to steric restraints on RNase I access (for a schematic overview see Figure S5A,B). Overall, biological replicates gave largely equivalent distribution patterns for both, yeast and human 80S FPs (Figure 1B,C and Figure S4). Collectively, these data validate the TCP-seq protocols used here, by presenting observations with 80S FPs that are largely equivalent to conventional Ribo-seq.

Next, we inspected human 40S FPs, which map throughout the 5’ region of mRNA including the initiation codon, while some are also found in the CDS (Figure 2A, see Figure S6A for replicate data). The latter show a length distribution indicating that they mostly derive from a small proportion of incompletely crosslinked 80Ses that did not withstand purification. These fragmented complexes produce a higher proportion of the smaller FP-type than seen in the 80S FP libraries (Figure S7A,B), consistent with a lower crosslinking efficiency for 80Ses with an empty A site. The 40S FPs also accumulate at stop codons (Figure S2A, right-handed panel), likely representing authentic intermediates of staged 80S recycling, wherein the 60S subunit (60S) is recycled to leave a 40S post-termination complex for the second stage of recycling (Hellen, 2018). Of note, 5’UTR 40S coverage, which should represent primarily scanning PICs, extends up to the mRNA 5’ ends (Figure 2B and Figure S7F). The broad size range of these 5’UTR 40S FPs, protecting mRNA fragments from a size corresponding approximately to the span of the 40S alone, of ∼26-nt, up to ∼70 nt (Figure 2C and Figure S8A), and their variable size distribution between replicates, is likely due to the transient nature of interactions within scanning PICs. We posit that this creates variability in the crosslinking efficiency, resulting in changes in RNase I access to cleavage sites not only at the boundaries but also within these assemblies. Together, these 40S FP distributions are consistent with the view that human 40Ses first bind mRNAs near the 5’ cap structure and then scan the 5’UTR as a larger assembly with the assistance of eIFs. All these observed patterns are similar to those seen in yeast (Figure S6B-D, S7C-E and G-I, and S8B-D) (Archer et al., 2016).

**Figure 2.**
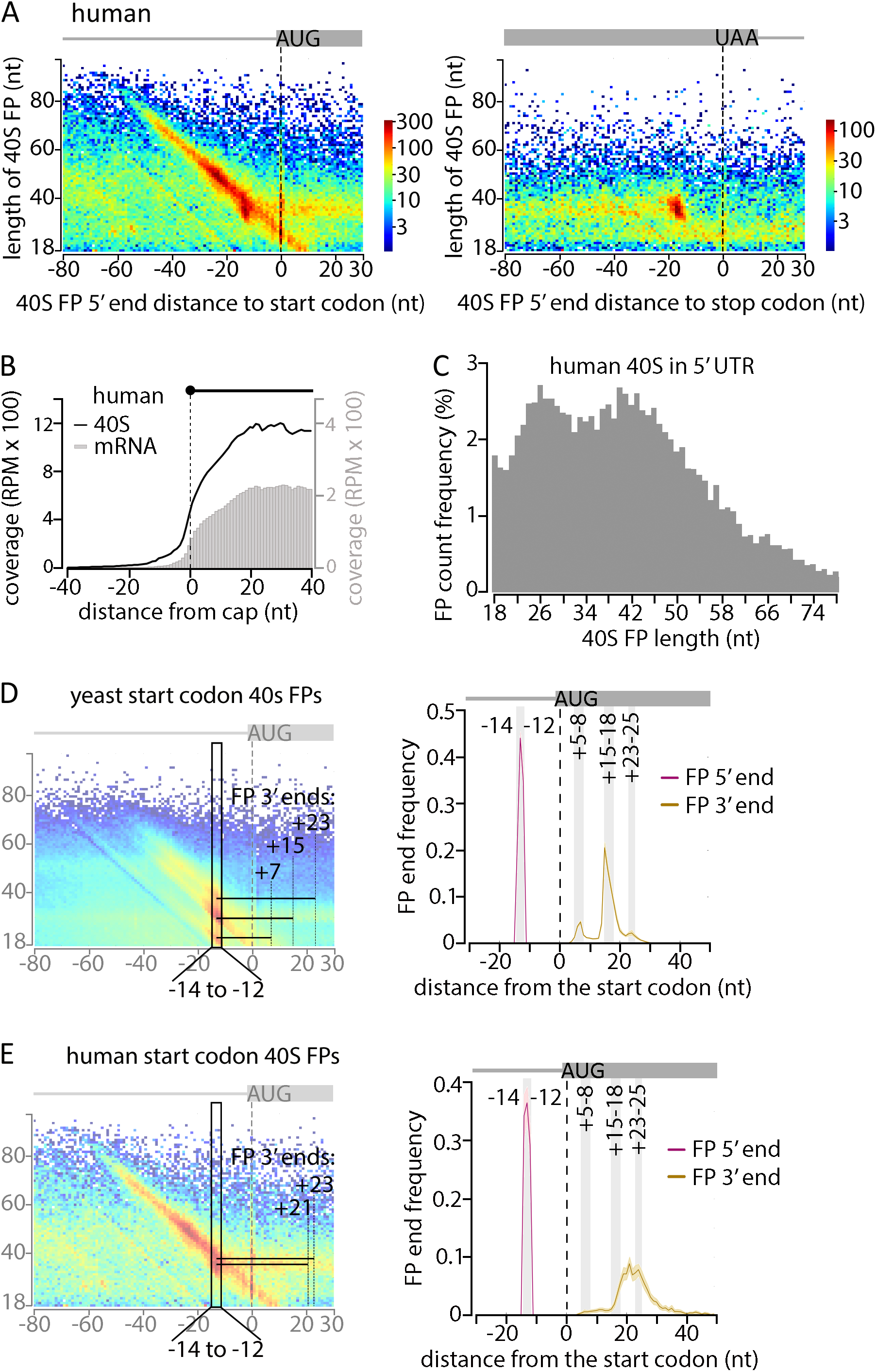
TCP-seq in human cells – small ribosomal subunit footprints. (A) Metagene plot of human 40S FP 5’ ends *versus* FP length. 40S FP 5’ ends are given relative to the first nucleotide (position 0) of start (left) or stop codons (right). Colour bars to the right of each plot indicate FP count in log10 scale. Data from 10,778 start sites and 16,803 stop sites is shown. (B) Metagene plot of human 40S FP coverage aligned to the annotated transcript start (cap). mRNA coverage (‘regular’ RNA-seq) is shown for comparison. Data from 136 mRNAs is shown whose annotated transcript start was confirmed (see STAR Methods) and whose annotated 5’ UTR has a minimal length of 80 nt, to avoid interference by the coverage of the 5’ extended start codon associated FPs. RPM, reads per million. (C) Length distribution of human 40S FPs assigned to the 5’UTR (excluding FPs mapped to the start codon region). Data from 9,241 genes are shown. (D) Metagene plot of 5’ and 3’ end position distributions of yeast 40S FPs that aligned to the start codon region of 5,994 mRNAs. FPs were chosen whose 5’ end map within the −14 to −12 positions relative to the start codon, as illustrated by the black rectangle in the heatmap on the left. Grey vertical bars in the distribution plot on the right indicate locations of discernible peaks. Those peaks are illustrated by representative FPs in the heatmaps on the left. The error regions shown as shading of the same colour indicate the 95^th^ percentiles of 1,000 iterations of random gene-wise resampling across all represented genes. (E) As (D) but for human 40S FPs. For comparison, vertical grey bars are shown with positions from (D). Discernable peaks in the 3’ end distribution on the right are illustrated by representative FPs in the heatmaps on the left. Data from 11,174 start sites are represented.

The original TCP-seq protocol had yielded three main yeast 40S FPs at start codons. Their 5’ ends predominantly mapped to position −12 (the A of the AUG is set to 0) with 3’ ends mostly found at +6, +16 and +24 positions, representing sequential intermediates of AUG recognition (Archer et al., 2016). We broadly replicate this pattern here, with only minor changes in the preferred yeast 40S FP 5’ and 3’ end positions (at −13, +7, +15 and +24, respectively) and in the relative proportions between FP of different sizes, with +15 representing the major intermediate (Figure 2D, see Figure S8E,F for replicate data; Figure S5C and S6E for graphical explanation). Another characteristic with start codon-associated FPs is that some exhibit extended 5’ ends beyond the dominant −13 position (Figure S6B-E and S5A,B). As their 3’ ends are unchanged, we assume that these 5’ extended FPs also have the start codon set in the 40S P-site. As noted above, the 5’ extensions likely represent eIF interactions at the 40S exit channel, expected for a scanning PIC, that are still partly in place (for FPs with a 3’ end reaching +5 to +8 positions) or a second ‘queuing’ 40S extending protection upstream (for FPs with a 3’ end reaching +15 to +18 and even more pronounced for those with a 3’ end reaching +23 to +25).

Human initiating 40Ses give a somewhat different pattern. A major FP 5’ position at −12/-13 is also seen, however, the major 3’ end shifted to +21 position (or +18, depending on the replicate) compared to +15 in yeast (Figure 2E, see Figure S8G for replicate data). Whereas in yeast all three start codon-associated 40S FPs, representing the main AUG recognition intermediates, feature 5’ extensions of comparable intensity as mentioned above (Figure S6B-D), notably, in human only the latest intermediate, i.e. that with the FP 3’ end position at +23 to +25, displays a prominent 5’ extension (Figure 2A and Figure S6A). The absence of dominant 5’ extensions at start codon-associated 40S FPs representing earlier intermediates might point to biological differences in AUG recognition kinetics, such as a much faster conversion of human 40Ses from early to late AUG-recognition states and/or a much slower 60S joining to convert initiating 40Ses into full 80Ses.

Taken together, our yeast findings obtained with the revised TCP-seq (and their suggested interpretations) largely mirror those found in the original TCP-seq study (Archer et al., 2016), illustrating high reproducibility of this approach. Despite the considerable evolutionary conservation of the initiation mechanism among low and high eukaryotes, mammalian TCP-seq pointed at numerous differences between yeast and humans that probably reflect variations in number of involved proteins, composition of initiation factor complexes and the means of their interactions with 40Ses, as well as with each other (Hinnebusch, 2017; Shirokikh and Preiss, 2018; Valášek, 2012).

### Selective (Sel)-TCP-seq detects staged dissociation of eIFs from translation initiation complexes and resolves individual start codon recognition steps

Next, we sought to deploy TCP-seq to better understand the role of individual eIFs during initiation. As different steps of the process associate with particular 40S positions along the mRNA and the factor composition of particular PICs at these positions may vary, our rationale was to compare the FPs of 40Ses associated with a factor of interest (FOI::40S) with those of unselected 40Ses. Thus, we subjected RNase I-digested 40Ses, prepared exactly as per the TCP-seq protocol, to an immunoprecipitation step using anti-FOI antibodies, creating Sel-TCP-seq (Figure 1A).

Our first FOI was eIF3 since it is the most complex translation initiation factor implicated in promoting numerous reactions throughout the translation cycle (for review see (Valasek et al., 2017)). We used our robust human eIF3b co-immunoprecipitation protocol (Wagner et al., 2014; Wagner et al., 2016), which efficiently pulls down the entire eIF3 complex, as well as other eIFs and the 40S (Figure S9A,B). In yeast, we took advantage of the well-established affinity tag pull downs (Nielsen et al., 2006; Valášek et al., 2002), and expressed the genes encoding the Tif32 and Nip1 subunits of eIF3 with a C-terminal FLAG tag from plasmids in a strain deleted for the corresponding genomic wild type allele (Figure S9C-E). Analogously, we generated a strain expressing a FLAG-tagged β subunit of the trimeric eIF2 complex chosen as our second FOI because it resides at the interface surface of the 40S, as opposed to the solvent side of the 40S where the eIF3 a/c subunits reside (Llacer et al., 2015). We sequenced FOI::40S FPs and compared them with FPs of the unselected pool of 40Ses isolated from the same experiment. Reassuringly, we observe a high correlation between the FOI::40S and the total 40S FP count per gene for each targeted factor and organism (Figure 3A-C, Figure S10A). Further, eIF2::40S and eIF3a/c::40S FP coverage in yeast, and eIF3b::40S coverage in humans, begins immediately at the 5’ end of mRNAs (Figure 3D-F, Figure S10B). This provides direct evidence for the long-standing expectation that most, if not all, 40Ses have both factors bound upon mRNA recruitment.

**Figure 3.**
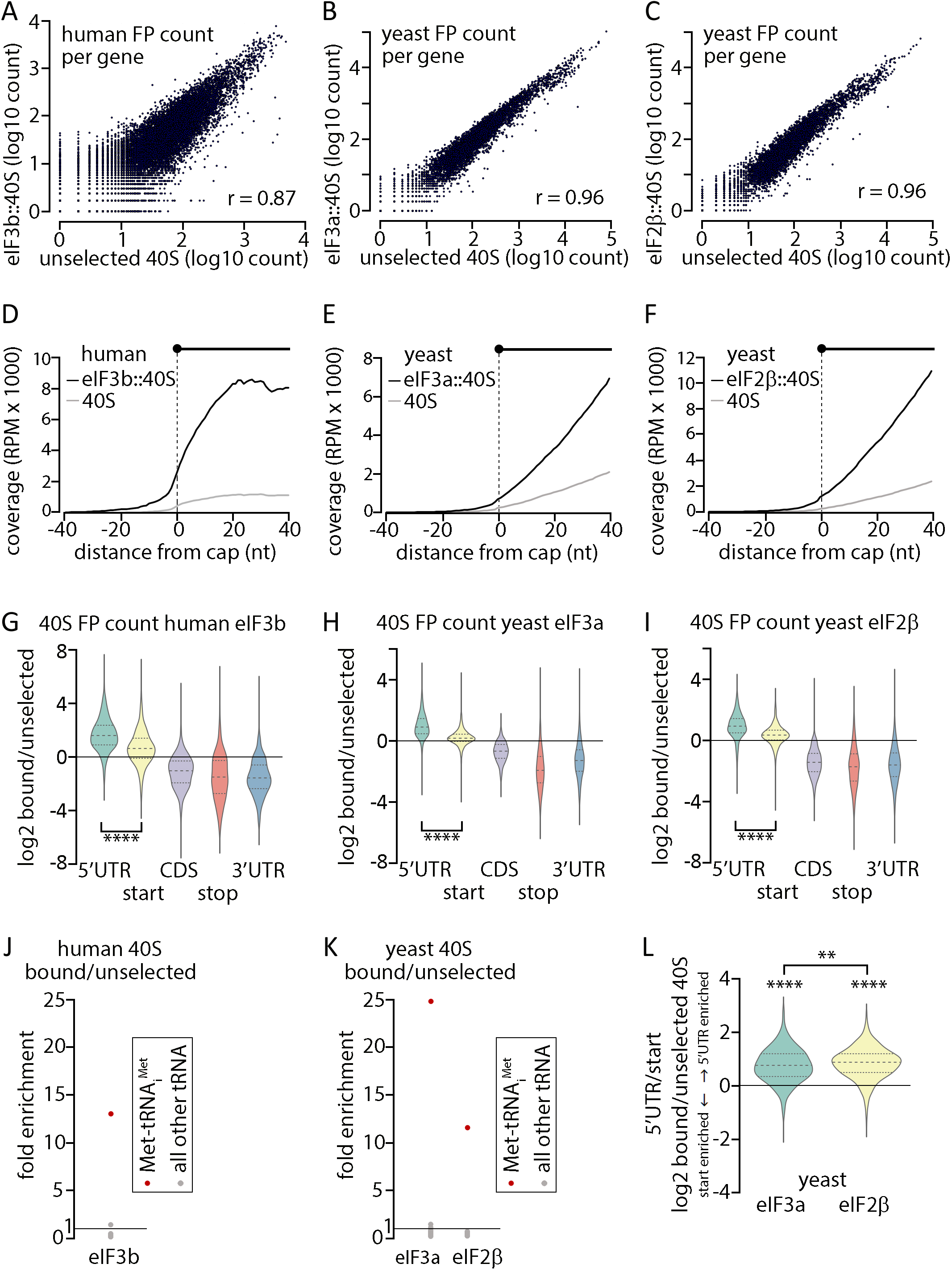
Sel-TCP-seq detects staged dissociation of eIFs from translation initiation complexes. (A) Scatter plot of the log10 count of human unselected 40S versus eIF3b::40S FPs mapped per gene. A pseudo count of 1 was added to each gene. r, Pearson correlation coefficient. (B) As (A) but for yeast unselected 40S FPs versus eIF3a::40S FPs mapped per gene. (C) As (A) but for yeast unselected 40S FPs versus eIF2β::40S FPs mapped per gene. (D) Metagene plot of human eIF3b::40S FP coverage aligned to the annotated transcript start (cap). Unselected 40S FP coverage is shown for comparison. Data from 136 mRNAs is shown whose annotated transcript start was confirmed (see STAR Methods) and whose annotated 5’ UTR has a minimal length of 80 nt, to avoid interference by the coverage of the 5’ extended start codon associated FPs. RPM, reads per million. (E) As (D) but for yeast eIF3a::40S and unselected 40S FP coverage. Data from 190 mRNAs are shown. (F) As (E) but for yeast eIF2β::40S and unselected 40S FP coverage. (G) Violin plots of human 40S FP counts assigned to the indicated transcript areas. Log2 ratios of eIF3b::40S versus unselected 40S FP counts are plotted. Genes with a minimal count of 100 per gene were considered (n = 1,189). A pseudo count of 1 was added to every transcript area. Two sample t-test was performed to calculate the indicated p-value (**** p << 0.0001). (H) As (G) but for yeast eIF3a::40S versus unselected 40S FPs (n = 2,142). (I) As (G) but for yeast eIF2β::40S versus unselected 40S FPs (n = 1,778). (J) Enrichment of tRNA fragment counts in the human eIF3b::40S versus the unselected 40S library. (K) Enrichment of tRNA fragment counts in the yeast eIF3a::40S library versus the unselected 40S library, and the yeast eIF2β::40S versus the unselected 40S library. (L) Violin plot of yeast 40S FP counts assigned to the 5’UTR versus those assigned to the start codon. The log2 of the FOI::40S FP count ratio vs. the unselected 40S FP count ratio is plotted. The relevant FOI is indicated below the x axis. Only genes with a minimal count of 10 in each of the transcript areas were considered (n = 1,053). One or Two sample t-tests were performed to calculate the indicated p-values (**** p << 0.0001, ** p < 0.01).

As detailed above, unselected 40S libraries contain a proportion of elongating 80S-derived FPs. They also feature fragments of both elongator tRNAs and Met-tRNA_i_^Met^, as seen before (Archer et al., 2016). For the most part, these CDS-located 40S complexes are not expected to involve either of the examined eIFs. Consistent with this, compared to unselected 40S FPs, eIF2::40S and each of the (yeast and human) eIF3::40S FPs are significantly underrepresented in the CDS and 3’UTRs but enriched in 5’UTRs (Figure 3G-I and Figure S10C), and their FP libraries are dramatically enriched for Met-tRNA_i_^Met^ fragments (Figure 3J,K and Figure S10D), demonstrating the selectivity of our approach. Importantly, we found that, compared to unselected 40S, eIF2::40S and each of the eIF3::40S are less enriched in the start codon region relative to the 5’UTR (Figure 3G-I and Figure S10C), confirming the expectation that both eIFs begin to dissociate from PICs during late stages of start codon recognition. Furthermore, the observation that this enrichment is significantly more pronounced for eIF2::40Ses than eIF3::40Ses in yeast (Figure 3L and Figure S10E) indicates that some start codon associated PICs still contain eIF3 but not eIF2. This is consistent with the generally accepted model that eIF2, together with eIF5, must leave the PIC upon AUG recognition, whereas a certain proportion of eIF3 complexes lingers on even during the early elongation cycles (for reviews see (Hinnebusch, 2017; Shirokikh and Preiss, 2018; Valasek et al., 2017)).

Comparing the FP 3’ end frequency of unselected start codon 40Ses with eIF2::40Ses or eIF3::40Ses in yeast reveals an enrichment of the longest 3’ end position (+23-25) at the expense of the shorter 3’ ends (mostly +15-18) for the FOI::40Ses (Figure 4A). Furthermore, the indicated enrichment is stronger for eIF3 versus eIF2. In view of our earlier interpretation that the start codon FP with the longest 3’ end represents a late PIC intermediate (Archer et al., 2016), these observations are consistent with the expectation that eIF2 leaves the initiating PIC before eIF3. The logic we apply to order unselected PIC intermediates represented by differing FP 3’ ends into a time sequence is based on the extent of evidence for a queuing 40S upstream of the initiating 40S, which was recently demonstrated (Shirokikh et al., 2019) and which is inferred from the 5’ extension of FPs, pronounced at a distance to fit a queuing 40S (around −30 nt) (see schematic in Figure S5A). These 5’ extensions are seen most prominently for the FPs with the longest 3’ end. With Sel-TCP-seq, a higher pull-down efficiency can be expected for PICs cross-linked to a queuing 40S, because the queuing 40S should contain both eIF2 and eIF3, whereas the initiating 40S might already have lost these factors. Assuming that late intermediates still have eIF3 bound while eIF2 has already dissociated, a higher pull down efficiency is expected for eIF3 compared to eIF2 because a higher proportion of PICs will have two epitopes of eIF3 (on the “late” and queuing 40Ses) but only one for eIF2 (on the queuing 40S) (for an illustration see Figure 4B).

**Figure 4.**
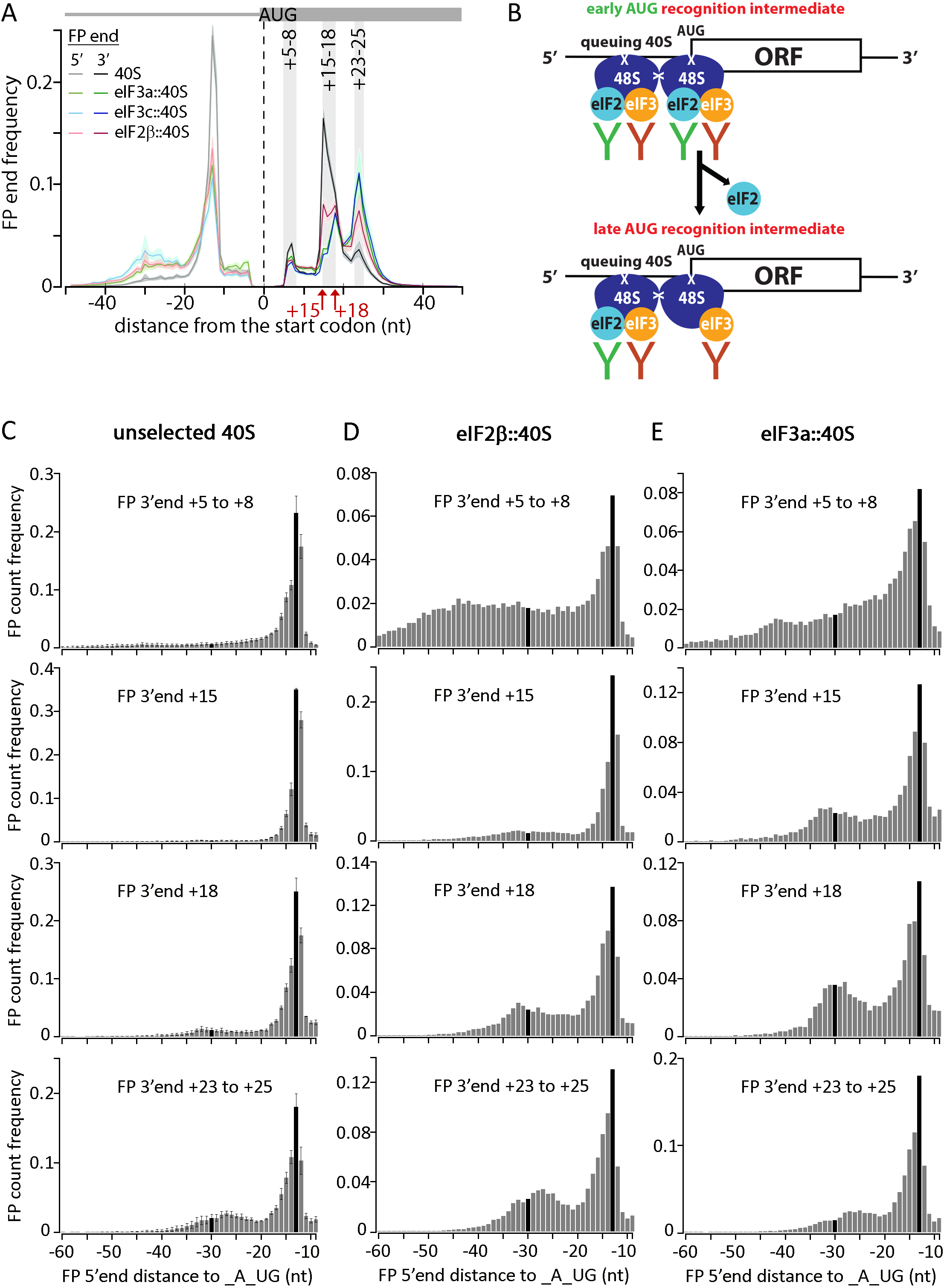
Sel-TCP-seq resolves individual start codon recognition steps. (A) Metagene plot of 5’ and 3’ end position distributions of yeast unselected and FOI bound 40S FPs that aligned to the start codon region of 5,994 mRNAs. Grey vertical bars indicating locations of discernible peaks are shown with positions from Figure 2D. The error regions shown as shading of the same colour indicate the 95^th^ percentiles of 1000 iterations of random gene-wise resampling across all represented genes. (B) Schematic of the suggested influence of queuing 40Ses on Co-IP efficiency. (C) 5’ end distribution of 40S FPs with specific 3’ end positions (relative to the A of the AUG start codon) as indicated in each panel. Selection matches the grey bars in (A) but respecting the partition of the +15 to +18 area into two peaks). For orientation, bars representing the −30 and −13 positions are highlighted in black. (D) As (C) but for eIF2β::40Ses. (E) As (C) but for eIF3a::40Ses.

Another interesting observation is the apparent split of a FP population with 3’ ends around +15-18 into two sub-populations, which are differently enriched with either eIF2::40S (+15 and +18) or eIF3::40S (primarily +18), suggesting that Sel-TCP-seq dissects the yeast AUG recognition process into four stages (Figure 4A). As stated above, to better interpret these distinct FP-types it is useful to group them by their 3’ end position and examine their extended 5’ ends (Figure 4C-E). Unselected 40S FPs then show a transition from a smooth trail up to around position −60 (representing scanning-type interactions) to an increasingly pronounced ‘bump’ at around −30 (representing a second, ‘queuing’ 40S), when they are sorted by increasing 3’ end positions (Figure 4C, top to bottom panels). We thus conclude that the FPs with their 3’ end at +15 represent the stage that follows the ‘+5-8 stage’ but precedes the ‘+18 stage’. Implying that the +15-18 set of FPs exemplifies PICs in the closed conformation, we propose that the +15 peak embodies 40Ses that have undergone the conformational switch from the open to closed state with the eIF2-TC still in the P^OUT^ arrangement, whereas the ‘+18 stage’ reflects the closed state with the TC already stabilized in the P^IN^ arrangement.

Closer inspection of the FP 5’ end distribution patterns of the +5-8 FP 3’ end position in the eIF2::40S and eIF3::40S *versus* unselected 40S libraries displays a marked enrichment of FPs with the longest 5’ end extensions (Figure 4C-E and Figure S10F, upper panels). Since eIF2 and eIF3 are both firmly established as scanning promoting factors (Hinnebusch, 2017), this finding supports our earlier proposal that the +5-8 stage represents PICs that are still partially in the scanning configuration (Archer et al., 2016). Strikingly, whereas the 5’ end distribution pattern of eIF3::40S FPs with 3’ ends at the +23-25 positions closely resembles that of the corresponding FPs of unselected 40Ses (Figure 4C and E, bottom panels), a distinct peak (at −32/33 FP 5’ end position) is revealed for those with 3’ ends at the +15 and +18 positions (Figure 4C and E, middle panels), whose nature can be explained as follows. As mentioned above, these two 3’ end positions likely represent the open-to-closed transition that is known to be accompanied by substantial conformational rearrangements (Llacer et al., 2015; Simonetti et al., 2016; Valasek et al., 2017). In particular, it has been suggested that the quaternary module of eIF3 (b-i-g-a–Cter.) relocates from the 40S solvent side (Figure S1C) to the subunit interface (Figure S1D) to contact eIF1 and eIF2γ at early stages of codon:anticodon recognition (represented by the “+15 peak”, Figure S1D) (Llácer et al., 2018). It was also proposed that after the completion of all necessary structural rearrangements stabilizing the TC in the P-site (possibly the “+18 peak”, Figure S1D), this quaternary module returns back to its original position (tentatively the “+23-25” peak, Figure S1C). Hence, we speculate that this temporal eIF3 relocation step around the entry channel allosterically provokes structural changes also around the exit channel, where the rest of eIF3 resides and which may involve also other eIFs, resulting in a transient protection of a larger piece of the exiting mRNA (creating the −32/33 5’ end peak, Figure S1D).

These findings demonstrate that deploying a selective approach to translation complex isolation (Sel-TCP-seq) enables a fine dissection of the individual initiation phases, as well as of the compositional/conformational states of PICs. Importantly, Sel-TCP-seq can help answering questions concerning functions of particular eIFs directly in the cell. It will be intriguing to employ Sel-TCP-seq in deciphering physiological roles of, for example, the mRNA-recruiting eIF4F complex and the subunit joining-promoting eIF5B factor.

### Sel-TCP-Seq reveals another layer of regulation of GCN4/ATF4 mRNAs and factor-dependence of the reinitiation-promoting short uORFs in yeast and human cells

The *ATF4* (in mammals) and *GCN4* (in yeast) genes encode master transcriptional activators and represent classical examples of genes under REI-mediated translational control in response to stress (Gunisova et al., 2018). The current model for the *GCN4* mechanism is illustrated in Figure S2A. Since the key tenets of this delicate mechanism were established mostly based on extensive yeast genetic analysis, we were curious to examine if they would find additional support in our TCP-seq data (Figure 5). As our experiments were done in nutrient-replete conditions, there was little detectable translation of the main *GCN4* CDS. Instead, unselected 40S and 80S coverage frequency is high across the 5’UTR of *GCN4* mRNA, broadly tracking uORF presence but with distinct local differences in the extent of their accumulation (Figure 5A, Figure S11A,B). Within the nAuORF1/2 region, 40Ses do not noticeably accumulate at the two potential non-AUG start codons and instead there is a broad and pronounced 40S peak in the region between them. As there are only modest levels of 80Ses throughout nAuORF1/2, these findings together indicate that initiation events are rare at both nAuORF1 and 2 start sites, consistent with the observation that they are dispensable for *GCN4* translational control (Zhang and Hinnebusch, 2011). Instead, most 40Ses appear to slow or stall in their scanning some distance into nAuORF1/2, at a site that overlaps with one of the REI-promoting elements linked to uORF1, RPE ii., positioned 69 nt downstream of the nAuORF1 start site, which forms a 9 base pairs-long hairpin (Figure 5B) (Gunisova et al., 2016; Munzarová et al., 2011). Altogether, this suggests that one of the main functions of the nAuORF region is exerted by the uORF1-linked RPE ii., which represents a barrier that scanning 40Ses have to overcome to reach the regulatory AUG-initiated uORFs. We propose that these stalled 40Ses might constitute a reservoir of initiation-competent (eIF2-TC containing) PICs, built up under normal growth conditions that can be utilized under stress conditions. Consistent with this, both eIF3::40S and eIF2::40S are enriched relative to unselected 40S in the nAuORF region, while this ratio inverts over uORFs 1-4 (Figure 5C,D). Combined with high abundance of *GCN4* mRNA (Hinnebusch, 2005), this then might assist with a fast and sustained Gcn4 upregulation, as it will be unaffected by the limiting amounts of the eIF2-TC responsible for the general translational shutdown during stress (Figure 5B).

**Figure 5.**
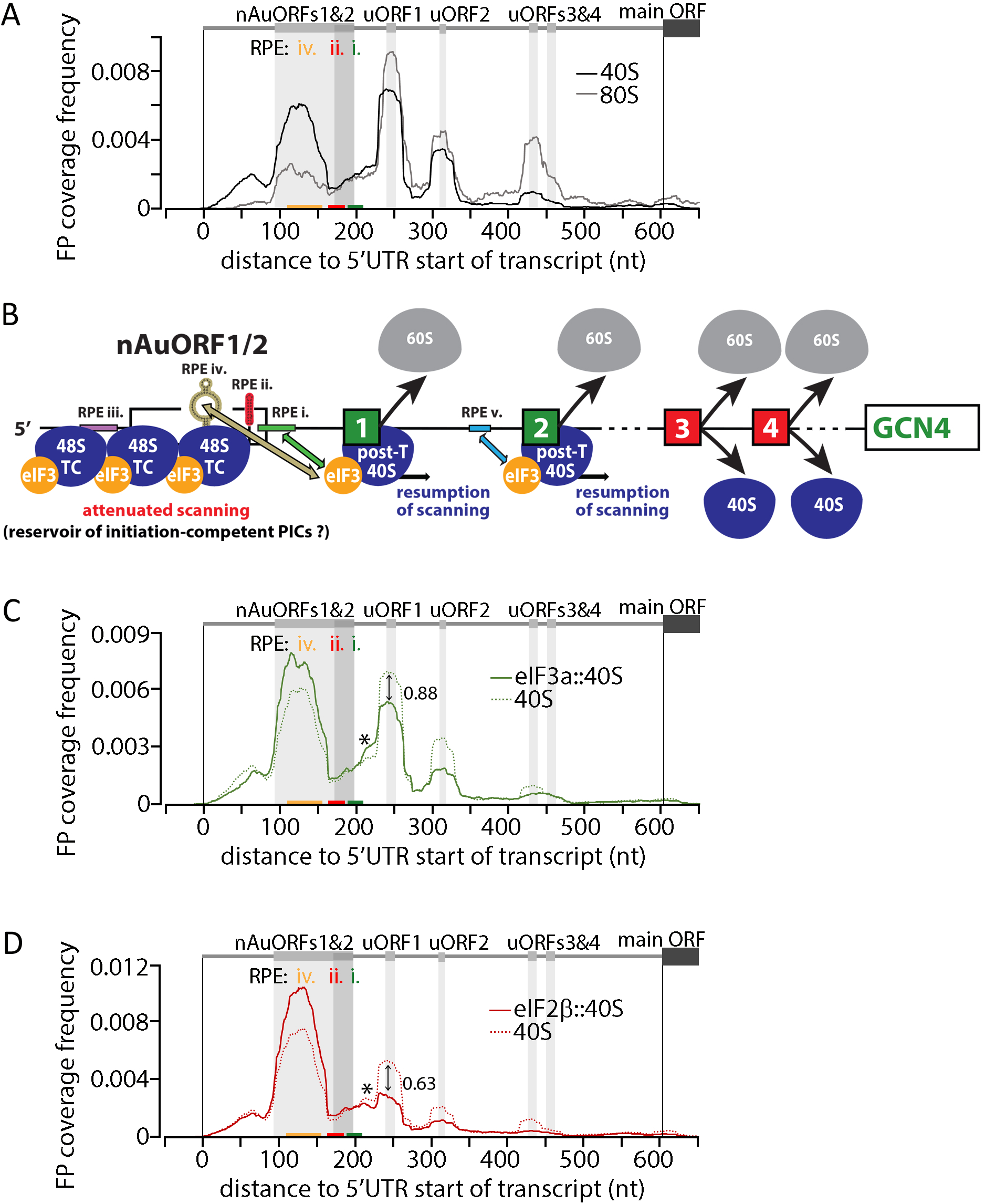
Sel-TCP-seq dissects reinitiation-mediated control of *GCN4* mRNA translation. (A) FP coverage frequency over the 5’UTR of *GCN4*. Separate tracks for yeast 40S and 80S FPs are shown. The location of AUG and non-AUG initiated uORFs are indicated with grey vertical bars. The location of the reinitiation promoting elements (RPE) i., ii., and iv. of uORF1 are indicated with colored horizontal bars. (B) Schematic of the yeast *GCN4* translational control mechanism. See text for further details. (C) As (A) but eIF3a selective 40S FP track and unselected 40S FP track is shown. The eIF3a::40S/unselected 40S ratio over uORF1 is given. The star indicates the “leading shoulder” described in the text. (D) As (C) but eIF2β selective 40S FP track and unselected 40S FP track is shown.

Interestingly, coverage in general is less for the second uORF in each pair (1 *vs.* 2 and 3 *vs.* 4), supporting the recently proposed “fail-safe” model of *GCN4* translational control postulating that uORF2 serves as the REI-promoting backup for uORF1, while uORF4 serves as an inhibitory backup for uORF3 (Gunisova and Valasek, 2014). Further, we noticed substantial differences in the 40S/80S ratios between the uORFs1,2 and uORFs3,4 pairs (Figure 5A, Figure S11A,B) (owing to their short length it is difficult to draw conclusions about uORF start or stop codon coverage *per se*). The relatively low 40S/80S ratios over uORFs3 and 4 can be explained by efficient conversion of 48S PICs into elongation-competent 80Ses followed by rapid termination/recycling for these uORFs. In contrast, the relatively high 40S/80S ratios over uORFs 1 and 2 might stem from a slow conversion of the post-termination 40Ses into those that resume 5’UTR traversal/scanning downstream. Why would that be? We recently proposed that the RPEs upstream of uORF1 (and uORF2) progressively fold into a specific secondary structure as the 40S scans through them (Munzarová et al., 2011). Following initiation at uORF1 and uORF2, eIF3 was shown to remain 80S-bound throughout the elongation and termination steps at these uORFs (Mohammad et al., 2017). Upon termination, eIF3 interacts with the folded RPEs i. and iv. to prevent 40S recycling specifically on these two REI-permissive uORFs (Gunisova et al., 2018). Hence, we speculate that these folding, binding and stabilization reactions might slow down transition through individual translation phases at the first two uORFs resulting in increased 40S FP occupancy, in contrast to the canonically initiated and terminated uORFs 3 and 4.

In this context, higher coverage frequency of eIF3::40S compared to eIF2::40S at uORF1 (0.88 over 0.63, relative to their respective unselected 40S tracks; Figure 5C,D and Figure S10G), as well as higher coverage frequency of eIF3::40S at uORF1 compared to uORF3 (0.88 over 0.59) could support the special eIF3 involvement in the REI-promoting mechanism of uORF1. However, a similar difference in coverage frequency of eIF3::40S compared to eIF2::40S is seen also with uORF3 (0.59 over 0.36), thus preventing us from drawing a clear conclusion. Nonetheless, note the selective occurrence of a coverage peak shoulder (due to the extended FP 5’ ends) preceding the uORF1 start codon in eIF3::40S *vs.* eIF2::40S (Figure 5C,D and Figure S10G). This might indicate that the presence of eIF3 (its interactions with RPEs; perhaps mainly with RPE i. immediately preceding the uORF1 start codon) is required for post-termination 40Ses in order to prevent their complete recycling, as proposed earlier based on yeast genetics and biochemical experiments (Figure 5B) (Gunisova et al., 2018).

Comparing the 40S and 80S coverage along the 5’UTR of *ATF4*, whose current regulatory model is illustrated in Figure S2B, reveals some similarities to the case of *GCN4*. Again, translation of the main *ATF4* CDS is very low and instead unselected 40S and 80S coverage broadly track each other along the 5’UTR but show distinct local differences. The 40S coverage frequency is high in a region overlapping with uORF0, accompanied by somewhat lower 80S coverage frequency but still higher than over the other two uORFs. This pattern resembles the high occupancies of 40Ses in the nAuORF1/2 region of *GCN4*, and could suggest a similar build-up of a reservoir of initiation-competent 40Ses upstream of the *ATF4* uORF1. However, the uORF0 proximity to the mRNA cap allowing for only one initiation-competent 40S to constitute this reservoir makes this option less likely. Interestingly, uORF0 has a special character because its “CDS” consists of just the AUG start codon immediately followed by the stop codon. Therefore, the 40S and 80S coverage frequency over this peculiar uORF might rather reflect the particular kinetics of non-canonical translation events, i.e. the balance between the 48S PIC conversion into the 80S initiating complex, its direct transition into the 80S termination complex, and emergence of the 40S post-termination recycling intermediate. As for the canonical *ATF4* uORFs, the higher 40S/80S ratio for uORF1 compared to uORF2 is broadly similar to that seen for uORF1,2 *versus* uORF3,4 on *GCN4 mRNA* (Figure 6A and Figure S11C). Importantly, we also detected an increased occupancy of heIF3::40Ses relative to unselected 40Ses at uORF1, when compared to uORF0 and uORF2 (Figure 6B). Our observations thus suggest that 1) the uORF0 region might form an active road block preventing uORF1 translation to keep the basal level of *ATF4* expression down (Figure 6C); and 2) the 40S reinitiation competence past uORF1 translation involves eIF3, as in case of yeast *GCN4*. Taken together, we conclude that the synthesis of these two functional orthologues is controlled by similar molecular mechanisms, despite the differences in the arrangement of their 5’UTRs.

**Figure 6.**
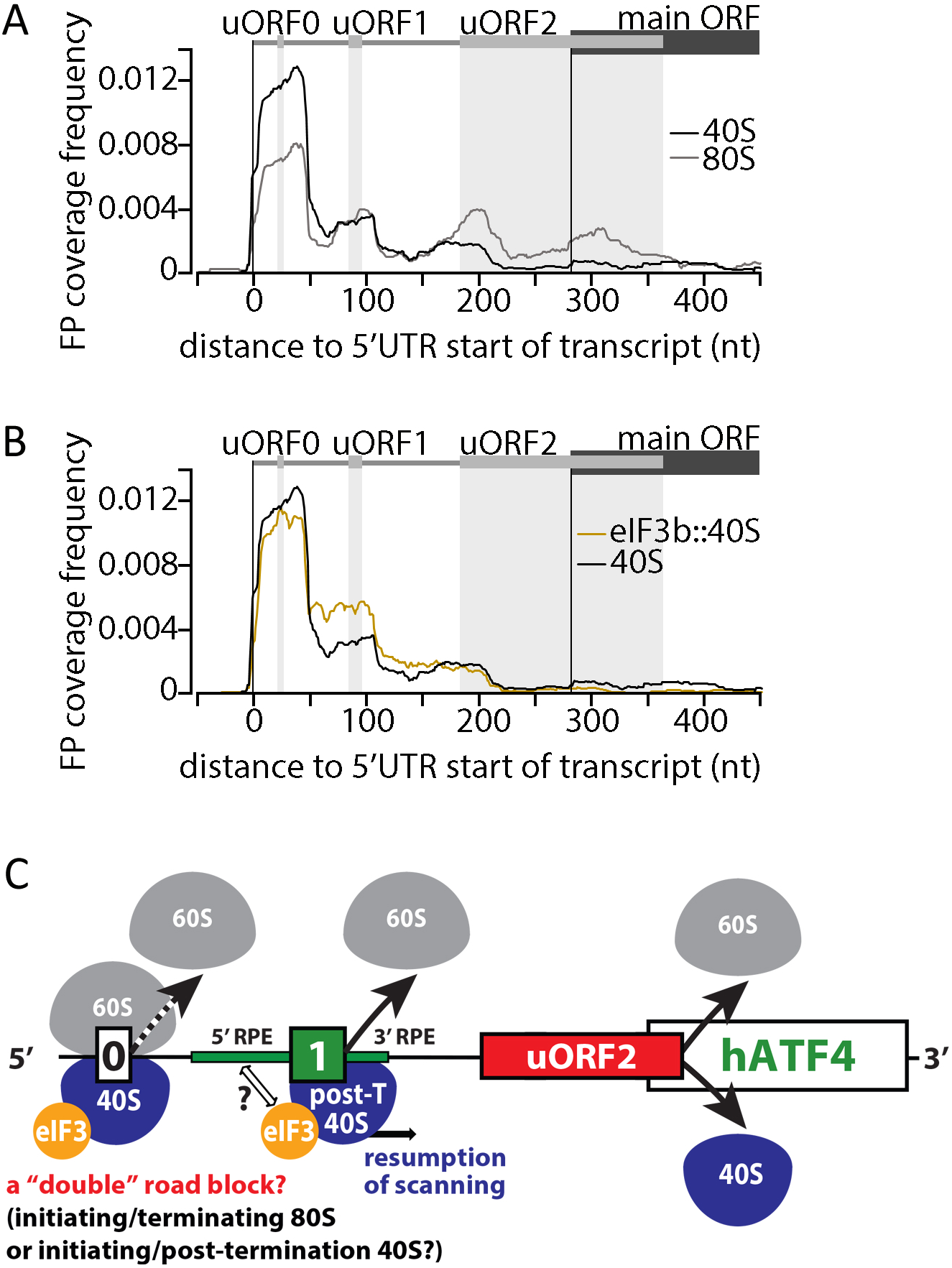
Sel-TCP-seq dissects reinitiation-mediated control of *ATF4* mRNA translation. (A) FP coverage frequency over the 5’UTR of *ATF4*. Separate tracks for human 40S and 80S FPs are shown. The location of uORFs 0, 1 and 2, is indicated with grey vertical bars. (B) As (A) but in addition to the 40S track the eIF3b selective 40S track is shown. (C) Schematic of the human *ATF4* translational control mechanism. See text for further details.

### Sel-TCP-seq enables monitoring of the co-translational assembly of higher order protein complexes within the cell

Sel-TCP-seq also affords a comparison of the FPs of 80Ses associated with a factor of interest to those of unselected 80Ses (Figure 1A). Translatome-wide analysis of yeast eIF3::80S FPs revealed dramatically uneven distribution within the CDS of several mRNAs encoding either non-targeted eIF3 subunits or other eIF3-associated eIFs. Closer inspection revealed that these patterns related to co-translational assembly of the targeted FOI with its interacting partner(s) under synthesis. In particular, we saw a sudden increase in 80S-FP coverage on the mRNA encoding the interacting partner at positions consistent with emergence of the nascent binding domain for the FOI from the ribosome exit tunnel (Figure 7A).

**Figure 7.**
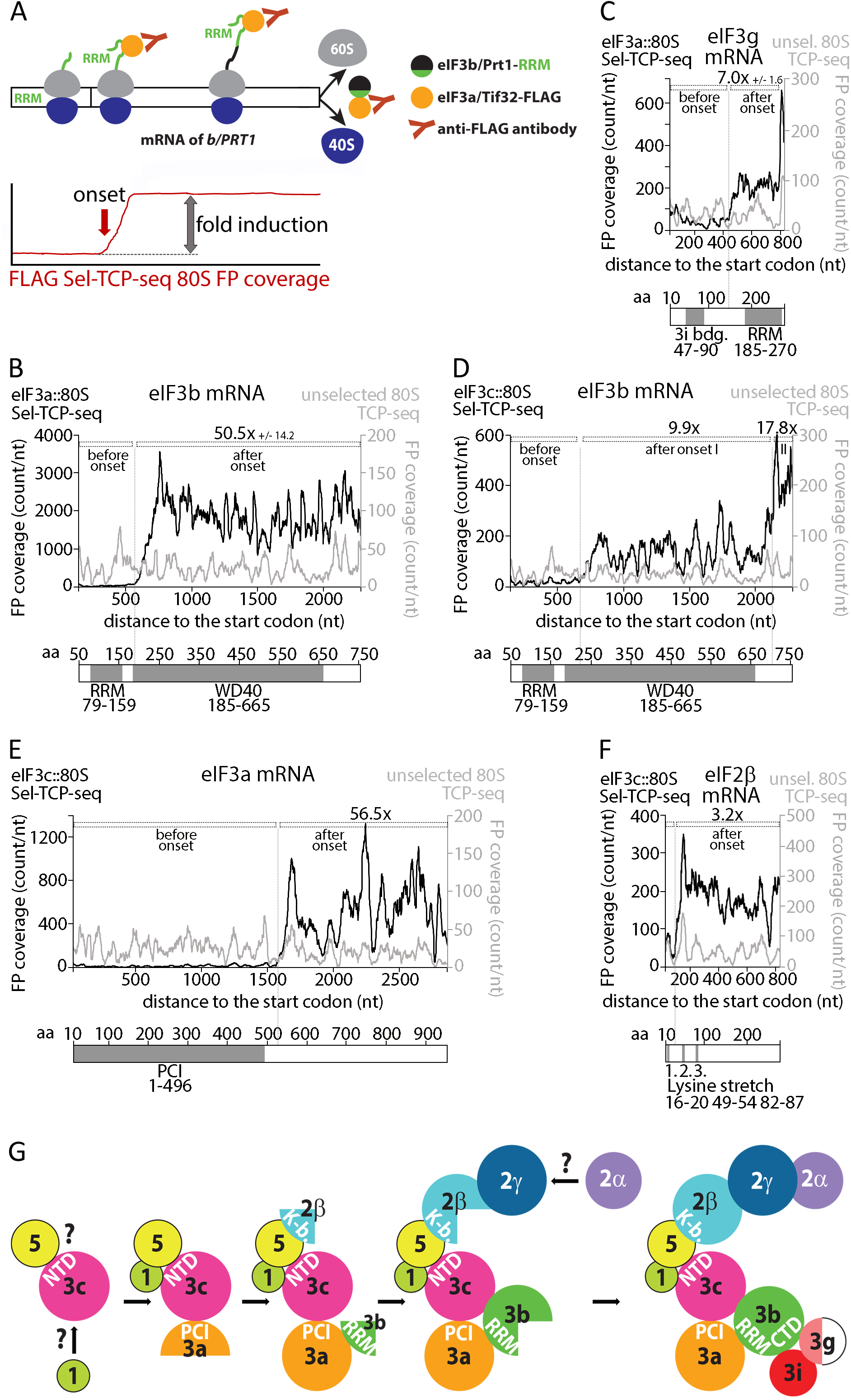
Sel-TCP-seq detects co-translational assembly within the multifactor complex in yeast. (A) Schematic of how co-translational interactions with a tagged FOI subunit lead to 80S-FP coverage increase within the CDS of mRNA encoding its assembly partner. The chosen example is the co-translational binding of nascent yeast eIF3b by eIF3a. RRM: RNA recognition motif. See text for further details. (B) -(F) Selected (black) and unselected 80S FP (grey) coverage tracks are shown for the following mRNAs encoding co-translationally interacting yeast proteins: eIF3b (B and D), eIF3g (C), eIF3a (E), eIF2β (F). The bar underneath the graphs depicts relevant known binding domains within the target subunit. FLAG-tagged targeted subunits are indicated in upper left corners. The coverage increase onset is indicated by the dotted line and the average fold induction of 3 replicates +/-SD is given for eIF3a-FLAG as bait. eIF3c-FLAG 80S Sel-TCP-seq was performed only once. Fold induction of the 80S Sel-TCP-seq coverage was normalized to that seen with unselected 80S TCP-seq (see STAR Methods for details). (G) The proposed *in vivo* MFC assembly model. Question marks indicate an unknown entry and form of the indicated factors into the assembly pathway. See text for details.

With yeast eIF3a as the FOI, we observed an ∼50 fold increase in the 80S FP coverage along the eIF3b (*PRT1)* mRNA for eIF3a::80S FPs versus unselected 80S FPs starting near nucleotide 580 and persisting until the end of the CDS (Figure 7B). The onset of this increase is ∼30 codons downstream of the CDS region that encodes the eIF3b RNA recognition motif (RRM; nucleotides 237 −477), which is one of the two main interacting domains for eIF3a within the eIF3 complex (Figure S1A) (Valášek et al., 2002; Zeman et al., 2019). As the ribosome exit tunnel accommodates ∼24 amino acids in extended conformation and ∼38 amino acids in α-helical conformation (Bhushan et al., 2010), the nascent RRM should have just emerged from the ribosome when an increase in FP coverage is seen. No eIF3a-mediated co-translational assembly was detected with eIF3c, the other well-established binding partner of eIF3a (see also below). However, we surprisingly detected evidence of co-assembly of eIF3a with eIF3g, as eIF3a::80S FP coverage increased ∼7-fold above unselected 80S FPs downstream of the eIF3g CDS region encoding its eIF3i-binding domain (Asano et al., 1998) (Figure 7C). Both eIF3g and eIF3i were previously shown to interact directly with the C-terminal region of eIF3b but not with eIF3a (Asano et al., 1998). Given the modest increase in eIF3a::80S FP coverage on eIF3g mRNA, and the fact that we did not detect any enhanced eIF3a::80S FP coverage within eIF3i mRNA, we posit that the co-translational assembly of eIF3g by eIF3a occurs indirectly, bridged by eIF3i already bound to eIF3b once the nascent eIF3i binding domain in eIF3g emerges from the exit tunnel on 80S translating eIF3g mRNA (Figure S12A; see also below).

Using yeast eIF3c as the FOI, we detected an increase in the eIF3c::80S FP coverage within the eIF3b mRNA just downstream of the CDS region encoding its RRM (Figure 7D), as noted above for the eIF3a::80S FPs. However, the increase for eIF3c::80S data was much less pronounced (∼10-fold; compared to ∼50-fold for eIF3a::80S), again suggesting that it is indirect, mediated by eIF3a, with which eIF3c directly interacts (Phan et al., 1998; Valášek et al., 2002). In accord with previous biochemical analyses, we detected a second, more robust increase in the eIF3c::80S FP coverage beyond the region coding for the eIF3b β-propeller domain (Figure 7D), and within the sequences encoding the eIF3b C-terminal domain, shown to interact directly with eIF3c (Valášek et al., 2002). Additionally, we see a very similar eIF3c::80S FP coverage pattern across the eIF3g mRNA CDS as described above for eIF3a::80S data, further suggesting that co-translational assembly of eIF3g must await association of the eIF3b/eIF3i heterodimer with eIF3a late in the pathway (compare Figure 7C and S13A).

Furthermore, the eIF3c::80S FP library revealed a strong (∼55-fold) increase in 80S FP coverage within eIF3a mRNA just downstream of the sequence encoding the PCI domain (Figure 7E), which is the only known eIF3c interaction domain within eIF3a (Figure S1A) (Valášek et al., 2002). The fact that we did not find evidence for co-translational assembly of eIF3c with eIF3a in the eIF3a::80S data (Figure S13B) suggests a uni-directional co-translational binding of the nascent PCI domain in eIF3a to fully synthesized eIF3c, but not *vice versa*. This type of uni-directional co-translational assembly was suggested to prevent increased propensity for aggregation/misfolding of proteins prone to these expression pathologies (Shiber et al., 2018).

Based on the above, we propose the following co-assembly pathway for yeast eIF3 (Figure S12B). The fully synthesized eIF3c co-translationally assembles with eIF3a forming an eIF3a-3c dimer, which subsequently picks up nascent eIF3b. At first, through a direct interaction between eIF3a and the eIF3b-RRM, later by an additional contact mediated by the helical region of eIF3c and the C-terminal end of eIF3b, forming a stable trimeric eIF3a-b-c subcomplex. This is consistent with earlier analysis demonstrating that this subcomplex is stable in cells, binds eIF1 as well as eIF5, and stimulates ribosome binding of mRNA and Met-tRNA_i_^Met^ *in vitro* (Phan et al., 2001)). Though, it contrasts with the human eIF3 assembly pathway, where the eIF3a and eIF3b subunits were proposed to form a nucleation core around which all other eIF3 subunits assemble into a 12-subunit complex (Wagner et al., 2016). This trimer then co-assembles with eIF3i but only once it is fully synthetized, contrary to eIF3g that can be picked up co-translationally. Targeting the remaining eIF3 subunits is required to examine whether there are alternative orders of co-assembly events; nonetheless, it is clear that eIF3 belongs to the growing class of multiprotein complexes that have been shown to undergo co-translational assembly by a ribosome profiling-based technique called SeRP (Shiber et al., 2018; Shieh et al., 2015).

Strikingly, the eIF3c::80S data also provides evidence for co-translational assembly of the β subunit of eIF2 with eIF3 soon after the first of three lysine stretches (dubbed K-boxes) of eIF2β emerges from the ribosome exit tunnel (Figure 7F). Since the eIF2β K-boxes were shown to interact selectively with either the ε subunit of eIF2B or the C-terminal domain of eIF5 (Asano et al., 2001b), but not with eIF3c (Asano et al., 2001b), this co-translational assembly is most likely again bridged by another factor. An indirect mode of co-translational assembly is also consistent with the modest ∼3-fold increase in eIF3c::80S FPs on eIF2β mRNA (Figure 7F). The prime candidate for the bridging factor is eIF5—a well-known interacting partner of eIF3c (Phan et al., 1998; Valášek et al., 2004) in addition to eIF2β, and a lynch pin in assembly of the MFC. Hence, we propose that eIF3c is the main nucleation core, not only of eIF3 but also of the entire MFC, fortifying the idea that eIFs 1, 3, 5, and eIF2-TC preassemble prior to joining the 40S subunit in 43S PIC formation (Asano et al., 2001a) (Figure 7G).

Interestingly, the recent SeRP analysis of eIF2 revealed that eIF2γ co-translationally assembles with eIF2β (Shiber et al., 2018), like seen here for eIF3c. Because the onset of the eIF2γ-targeted 80S FP coverage increase mapped to the eIF2γ–binding domain in the last third of the eIF2β CDS, it seems that the eIF3c-eIF5 dimer co-translationally interacts with eIF2β prior to its binding to eIF2γ; i.e. prior to the assembly of eIF2 *per se*. Whether it is only a transient, perhaps stabilizing interaction before eIF2 assembles as a trimeric complex or whether the entire MFC, and perhaps other higher order protein complexes can assemble co-translationally – independent of the integrity of their individual multi-subunit components – remains to be addressed. In any case, our findings nicely correlate with a recent report demonstrating that translation factor mRNAs are localized to distinct granules within growing yeast cells where they are actively translated, in contrast to other messenger RNP granules, such as P-bodies and stress granules, containing translationally repressed mRNAs (Pizzinga et al., 2019).

Overall, Sel-TCP-seq appears as a versatile and powerful technique for co-assembly discoveries, because the third-party-bridged, co-translational interactions that we observed here were not detected with previous approaches (Shiber et al., 2018). We speculate that it is the initial, rapid formaldehyde *in vivo* cross-linking step that stabilizes multiprotein assemblies well enough to detect even co-translational assemblies that are bridged by one or more proteins, which are characterized by a less robust 80S FP coverage increase than that seen for direct co-assemblies.

## CONCLUDING REMARKS

By modifying the yeast TCP-seq and adopting it for human cell lines, we revealed that although eukaryotic protein synthesis is evolutionary conserved, particular differences exist, such as in a factor occupancy of PICs at various initiation stages and in the AUG recognition kinetics, perhaps reflecting an increased complexity of translational control in higher eukaryotes. Furthermore, we demonstrated that the Sel-TCP-seq technique presented in this study provides researchers with a unique opportunity to dive deep not only into molecular details of assembly and general functioning of mRNA-associated ribosomal complexes *in vivo*, but also into regulatory mechanisms governing the actual translational output. In other words, with this technique we can now ask when, and where along the mRNA 5’UTR, interactions among eIFs (beyond those investigated here), other auxiliary factors, mRNA and ribosomal subunits are made and broken during individual phases of translation, and how these processes are regulated in response to changing environmental conditions.

## ACKNOWLEDGEMENTS

We are grateful to Nicolas Ingolia for sequencing and analysis of our first Sel-TCP-seq libraries as well as mentoring mainly at the onset of this study, and to the members of Valasek, Preiss, and Hinnebusch labs for helpful suggestions. We are also indebted to Olga Krýdová and Veronika Kovaničová for technical and administrative assistance. This work was supported by a Grant of Excellence in Basic Research (EXPRO 2019) provided by the Czech Science Foundation (19-25821X to L.S.V.), a Wellcome trust grant (090812*/*B*/*09*/*Z to L.S.V.), an Australian Research Council Discovery Project Grant (DP180100111 to T.P. and N.E.S), a National Health and Medical Research Council Principal Research Fellowship (APP116999 to R.D.H.) and a Cancer Council ACT Project Grant (APP1120469 to T.P. and R.D.H.).

## AUTHOR CONTRIBUTIONS

S.W., T.P. and L.S.V. conceived and designed the project.

S.W. carried out all experiments with collaborative contributions from A.H., V.H., N.S., and N.E.S..

S.W. performed the data analysis.

S.W., T.P. and L.S.V. interpreted results and wrote the paper with input from A.H., N.E.S., R.D.H. and A.G.H..

## SUPPLEMENTAL FIGURE LEGENDS

**Figure S1. Schematics of eIF3.** (A, B) In *S. cerevisiae*, the eIF3 complex is formed by 5 subunits (A), whereas in mammals it consists of 12 subunits (B). These schematics illustrate similarities and differences between budding yeast and mammalian eIF3. One of the main structural domains shared by several eIF3 subunits – the PCI (for Proteasome, COP9, Initiation factor 3) domain – is shown in bold in both panels; adopted from (Zeman et al., 2019). (C, D) Schematic representation of the arrangement of eIF3 subunits in the two available conformations that are deduced from the available Cryo-EM analysis (adapted from (Valasek et al., 2017)). (A) eIF3 residing exclusively on the solvent-exposed side with the octamer occupying the platform of the small ribosomal subunit connected with the eIF3b–g–i module – sitting near the mRNA entry channel – *via* the extended C-terminal linker domain of eIF3a (a dashed red line indicates a predicted location of the eIF3a-Cterm). The 40S subunit is depicted in grey surface; all other subunits are labelled and colored variably. The predicted path of mRNA is shown in dark red; Ex and En – mRNA exit and entry channels, respectively. This conformation of eIF3 likely generates FPs with 3’ end at +5 to +8 and +23 to +25 (see Figure 4E and text for details). (B) In this conformation, the entire eIF3a-Cterm–b–g–i module relocates from the solvent-exposed side to the intersubunit side, so that the eIF3b-RRM interacts with 18S rRNA and eIF1 and the eIF3b-propeller interacts with eIF2γ. This conformation of eIF3 likely generates FPs with 3’ end at +15 and +18 with a specific −32/33 5’ end peak (see Figure 4E and text for details).

**Figure S2. Translational regulation of GCN4/ATF4.**

(A) Schematic of *GCN4* 5’UTR and regulation. The REI-permissive uORF1 is translated under both nutrient-replete and depleted conditions. After its translation, only the 60S subunit is recycled, whereas the 40S subunit remains bound to the *GCN4* mRNA – due to a specific interaction between RPEs of uORF1 and eIF3 (Mohammad et al., 2017; Munzarová et al., 2011) – to eventually resume traversing downstream. It cannot start “scanning” *per se* for the next AUG until it re-acquires an active eIF2-TC, the levels of which are reduced under starvation/stress conditions. uORF2, functionally mimicking uORF1, was proposed to serve as its backup to capture 40Ses that ‘leaky-scanned’ past uORF1, thereby maximizing the REI potential as a fail-safe mechanism (Gunisova and Valasek, 2014). In non-starved cells, where eIF2-TC levels are high, nearly all of the 5’ to 3’-traversing 40Ses rebind the eIF2-TC before reaching the REI-non-permissive uORF3 or uORF4. Upon their translation, terminating ribosomes undergo full ribosomal recycling, thus preventing REI at the main CDS of *GCN4.* It is noteworthy that uORF4 and not the preceding uORF3 has been believed to represent the main negative element of this system (Hinnebusch, 2005) (see further below). When eIF2-TC levels are low, a large proportion of the 40Ss traversing downstream will bypass uORFs 3 and 4 and – upon eventual acquisition of the eIF2-TC – reinitiate at the *GCN4* start codon. Hence, whereas the stress response shuts down general translation initiation, as eIF2-TCs are required for translation of most mRNAs, it at the same time stimulates *GCN4* translation to trigger stress adaptation programs (reviewed in (Gunisova et al., 2018; Hinnebusch, 2005).

(B) Schematic of *ATF4* 5’UTR and regulation. The REI-permissive uORF1 is translated under both non-stress and stress conditions. Upon termination, the 40S remains bound to the *ATF4* mRNA to eventually resume traversing downstream. Under high levels of the eIF2-TC, 40Ses reinitiate at uORF2 whose translation prevents reinitiation at the *ATF4* start codon. Under low levels of the eIF2-TC, majority of 40Ses skip uORF2 and reinitiate at *ATF4* instead (Lu et al., 2004; Vattem and Wek, 2004). The role of uORF0 is unknown.

**Figure S3. Crosslinking and RNase I conditions. Replicate correlation.**

(A) Polysome profile of HEK293T cells either treated with cycloheximide (CHX) for 1 min (according to (Wagner et al., 2014)) or with different concentrations of formaldehyde (HCHO) according to the cross linking conditions outlined in the STAR Methods.

(B) Polysome profiles of whole cell extracts of formaldehyde cross-linked HEK293T cells digested with different amounts of RNase I.

(C) Scatter plot of the log10 count of human unselected 80S FPs mapped per gene of replicate 1 versus replicate 2. The counts were normalised to the library size. A pseudo count of 1 was added to each gene. r, Pearson correlation coefficient.

(D) As (C) for human 40S FPs.

(E) As (C) for yeast 80S FPs. Replicate 1-1 vs. 1-2.

(F) As (C) for yeast 80S FPs. Replicate 1-3 vs. 1-2.

(G) As (C) for yeast 80S FPs. Replicate 1-1 vs. 1-3.

(H) As (C) for yeast 40S FPs. Replicate 1-1 vs. 1-2.

(I) As (C) for yeast 40S FPs. Replicate 1-1 vs. 1-3.

(J) As (C) for yeast 40S FPs. Replicate 1-3 vs. 1-2.

**Figure S4. TCP-seq in yeast and human cells – full ribosome footprints. (Corresponding to** Figure 1)

(A) Metagene plot of replicate 2 of human 80S FP 5’ ends *versus* FP length. 80S FP 5’ ends are given relative to the first nucleotide (position 0) of start (left) or stop codons (right). Colour bars to the right of each plot indicate FP count in log10 scale. Data from 10,778 start sites and 16,803 stop sites are shown.

(B) As (A) but for yeast replicate 1-1. Data from 6,639 start sites and 6,514 stop sites is shown.

(C) As (B) but for yeast replicate 1-2.

(D) As (B) but for yeast replicate 1-3.

(E) Length distribution of replicate 2 of human 80S FPs assigned to annotated CDS (excluding start and stop codon associated FPs) of 9,241 genes is shown.

(F) As (E) but for yeast replicate 1-1. Data of 5,818 genes are shown.

(G) As (F) but for yeast replicate 1-2.

(H) As (F) but for yeast replicate 1-3.

**Figure S5. Schematics to facilitate read out of the heatmaps.**

Schematics facilitating the interpretation of the heatmaps displayed in Figures 1, 2, S4, S6. The schematics are hypothetical and depict the three main pattern we observed around the start codon.

(A) Illustration of a start codon-associated 40S/80S with a queuing 40S upstream and the resulting heatmap pattern of the footprints such an event would produce.

(B) Illustration of a start codon-associated 40S/80S with a line up of several other factors upstream and the resulting heatmap pattern of the footprints such an event would produce.

(C) Illustration of three start codon-associated 40S/80S protecting varying stretches of mRNA on the entry channel side (3’ end) and the resulting heatmap pattern of the footprints such an event would produce.

**Figure S6. TCP-seq in yeast and human cells – small ribosomal subunit −1. (Corresponding to** Figure 2A)

(A) Metagene plot of replicate 2 of human 40S FP 5’ ends *versus* FP length. 40S FP 5’ ends are given relative to the first nucleotide (position 0) of start (left) or stop codons (right). Colour bars to the right of each plot indicate FP count in log10 scale. Data from 10,778 start sites and 16,803 stop sites are shown.

(B) As (A) but for yeast replicate 1-1. Data from 6,639 start sites and 6,514 stop sites is shown.

(C) As (B) but for yeast replicate 1-2.

(D) As (B) but for yeast replicate 1-3.

(E) Schematic explaining the distinct 3’ ends of start codon-associated FPs.

**Figure S7. TCP-seq in yeast and human cells – small ribosomal subunit −2. (Corresponding to** Figure 2B)

(A) Length distribution of human 40S FPs assigned to annotated CDS (excluding start and stop codon associated FPs) of 9241 genes is shown. For replicate 1.

(B) As (A) but for human replicate 2.

(C) As (A) but for yeast replicate 1-1. Data from 5818 genes are shown.

(D) As (C) but for yeast replicate 1-2.

(E) As (C) but for yeast replicate 1-3.

(F) Metagene plot of human 40S FP coverage aligned to the annotated transcript start. mRNA coverage (‘regular’ RNA-seq) is shown for comparison. Data from 136 mRNAs are shown whose annotated transcript start was confirmed (see STAR Methods) and whose annotated 5’ UTR has a minimal length of 80 nt, to avoid interference by the coverage of the 5’ extended start codon associated FPs. RPM, reads per million. Replicate 2.

(G) As (F) but for yeast replicate 1-1. Data from 190 mRNAs are shown.

(H) As (G) but for yeast replicate 1-2.

(I) As (G) but for yeast replicate 1-3.

**Figure S8. TCP-seq in yeast and human cells – small ribosomal subunit −3. (Corresponding to** Figure 2C-E)

(A) Length distribution of replicate 2 of human 40S FPs assigned to the 5’UTR (excluding FPs mapped to the start codon region). Data from 9,241 genes are shown.

(B) As (A) but for yeast replicate 1-1. Data from 5818 genes are shown.

(C) As (B) but for yeast replicate 1-2.

(D) As (B) but for yeast replicate 1-3.

(E) Metagene plot of 5’ and 3’ end position distributions of yeast 40S FPs that aligned to the start codon region of 5,994 mRNAs. FPs were chosen whose 5’ end map within the −14 to −12 positions relative to the start codon. Grey vertical bars indicate locations of discernible peaks. The error regions shown as shading of the same colour indicate the 95^th^ percentiles of 1,000 iterations of random gene-wise resampling across all represented genes. Replicate 1-2 of corresponding Figure 2D.

(F) As (E) but for yeast replicate 1-3.

(G) As (E) but for human replicate 2 of corresponding Figure 2E. For comparison, vertical grey bars are shown with positions from yeast (Figure 2D, S8E,F). Data from 11,174 start sites are represented.

**Figure S9. Background controls for CoIPs.**

(A) Western blot of an anti-eIF3b co-immunoprecipitation from the 40S fractions of formaldehyde-crosslinked and RNase I-digested HEK293T cells.

(B) RNA gel of a anti-eIF3b co-immunoprecipitation from the 40S fractions of formaldehyde-crosslinked and RNase-I digested HEK293T cells.

(C) Western blot of an anti-FLAG co-immunoprecipitation from the 40S fractions of formaldehyde-crosslinked and RNase I-digested yeast cells expressing FLAG-tagged eIF3a.

(D) Re-sedimentation of the eluates of an anti-FLAG co-immunoprecipitation from the 40S fractions of formaldehyde-crosslinked and RNase I-digested yeast cells expressing FLAG-tagged eIF3a or untagged eIF3a.

(E) RNA gel of an anti-FLAG co-immunoprecipitation from the 40S fractions of formaldehyde-crosslinked and RNase I-digested yeast cells expressing FLAG-tagged eIF3a.

**Figure S10. Yeast eIF3c::40S Sel-TCP-seq.**

(A) Scatter plot of the log10 count of yeast unselected 40S versus eIF3c::40S FPs mapped per gene. A pseudo count of 1 was added to each gene. r, Pearson correlation coefficient. Corresponding to Figures 3B and C for yeast eIF3a::40S and eIF2β::40S, respectively.

(B) Metagene plot of yeast eIF3c::40S FP coverage aligned to the annotated transcript start (cap). Unselected 40S FP coverage is shown for comparison. Data from 190 mRNAs are shown whose annotated transcript start was confirmed by us (see STAR Methods) and whose annotated 5’ UTR has a minimal length of 80 nt, to avoid interference by the coverage of the 5’ extended start codon associated FPs. RPM, reads per million. Corresponding to Figures 3E and F for yeast eIF3a::40S and eIF2β::40S, respectively.

(C) Violin plots of yeast 40S FP counts assigned to the indicated transcript areas. Log2 ratios of eIF3c::40S versus unselected 40S FP counts are plotted. Genes with a minimal count of 100 per gene were considered (n = 1,536). A pseudo count of 1 was added to every transcript area. Two sample t-test was performed to calculate the indicated p-value (**** p << 0.0001). Corresponding to Figures 3H and I for yeast eIF3a::40S and eIF2β::40S, respectively.

(D) Enrichment of tRNA fragment counts in the yeast eIF3c::40S library versus the unselected 40S library. Corresponding to Figures 3K for yeast eIF3a::40S and eIF2β::40S.

(E) Violin plot of yeast 40S FP counts assigned to the 5’UTR versus those assigned to the start codon. The log2 of the FOI::40S FP count ratio vs. the unselected 40S FP count ratio is plotted. The relevant FOI is indicated below the x axis. Only genes with a minimal count of 10 in each of the transcript areas were considered (n = 1,053). One or Two sample t-test was performed to calculate the indicated p-values (**** p << 0.0001). Corresponding to Figure 3L comparing yeast eIF3a::40S and eIF2ẞ::40S.

(F) 5’ end distribution of 40S FPs with specific 3’ end positions (relative to the A of the AUG start codon) as indicated in each panel. Selection matches the grey bars in (A) but respecting the partition of the +15 to +18 area into two peaks). For better orientation, bars representing the −30 and −13 positions are highlighted in black. Corresponding to Figures 4D and E for yeast eIF2β::SSUs and eIF3a::SSUs, respectively.

(G) FP coverage frequency over the 5’ UTR of *GCN4*. Separate tracks for unselected SSU and eIF3c::40S FPs are shown. The location of AUG and non-AUG initiated uORFs are indicated with grey vertical bars. The location of the reinitiation-promoting elements (RPE) i., ii., and iv. of uORF1 are indicated with coloured horizontal bars. The eIF3c::40S/unselected 40S ratio over uORF1 is given. The star indicates the shoulder described in the text. Corresponding to Figures 5C and D for yeast eIF3a::40S and eIF2β::40S, respectively.

**Figure S11. Replicates for Figures 5 and 6.**

(A) FP coverage frequency over the 5’ UTR of GCN4. Tracks for yeast 40S FPs and 80S FPs are shown. The location of AUG and non-AUG uORFs are indicated with grey vertical bars. Replicate 1-2 of corresponding Figure 5A.

(B) As (A) but for replicate 1-3.

(C) As (A) but human replicate 2 of corresponding Figure 6A.

**Figure S12. Schemes to facilitate interpretations of** Figure 7.

(A) Schematic illustrating the co-translational interactions of the yeast eIF3a-b-i subcomplex with nascent eIF3g, revealed by FLAG co-immunoprecipitation in a strain expressing FLAG tagged eIF3a. RRM: RNA recognition motif.

(B) Schematic illustrating the co-translational assembly pathway of eIF3 in yeast. RRM: RNA recognition motif. NTD: N-terminal domain. CTD: C-terminal domain. WD40: WD40 domain. PCI: Proteasome-COP9-Initiation factor 3 domain.

**Figure S13. 80S-Sel-TCP-seq tracks for all MFC subunits; addition to** Figure 7.

(A) eIF3c-selected (black) and unselected 80S FP (grey) coverage tracks are shown for eIF3g mRNA. The bar underneath the graphs depicts relevant known binding domains within the target subunit. The coverage increase onset is indicated by the dotted. Fold induction of the 80S Sel-TCP-seq coverage was normalized to that seen with unselected 80S TCP-seq (see STAR Methods for details).

(B) eIF3a selected (upper row) and unselected (lower row) 80S FP coverage tracks are shown for all mRNA encoding eIF3 subunits.

(C) As (B) but for all mRNA encoding MFC subunits other than eIF3.

(D) eIF3c selected (upper row) and unselected (lower row) 80S FP coverage tracks are shown for all mRNA encoding eIF3 subunits.

(E) As (D) but for all mRNA encoding MFC subunits other than eIF3.

## STAR METHODS

Detailed methods are provided in the online version of this paper and include the following:

### LEAD CONTACT AND MATERIALS AVAILABILITY

Further information and requests for resources and reagents should be directed to and will be fulfilled by the Lead Contact Leoš Shivaya Valášek.

### SUBJECT DETAILS

#### Human cell line

The cell line HEK293T was used. Cells were grown in Dulbecco’s modified Eagle’s media supplemented with 10% fetal bovine serum at 37⁰C and 5% CO_2_. For experiments, 3.5 Mio cells in 20 ml of media were seeded into a Ø 15 cm dish and were grown for 48h before harvesting at approximately 80% confluency.

#### S. cerevisiae strains

Yeast strains SY182, SY183, LMY61 (SY194) and H25 were used in this study (Table S1). To generate SY182 and SY183, YBS52 ((Munzarová et al., 2011)) was transformed with pRS315-a/TIF32-FLAG-L (Klaus Nielsen) and Yep181-a/TIF32-L (Leoš Valášek), respectively, and YCp-a/TIF32-HIS-U was evicted by growth on 5-FOA. For experiments, yeast strains were grown at 30⁰C in 1 L of YPD up to an OD_600_ of 0.7-0.8.

### METHOD DETAILS

#### Formaldehyde cross-linking and preparation of whole cell extracts

**HEK293T —** Two Ø 15 cm dishes were processed together at any time. Dishes were transferred to a cold room and 600 µl of a 10 % (w/v) formaldehyde solution (freshly diluted in H_2_O at room temperature) were added to the media, resulting in a ∼0.3 % final concentration of formaldehyde. To accomplish fast and uniform mixing of the formaldehyde with the media, dishes were tilted and the formaldehyde solution was added into the pool of media, mixed, then dishes were turned back to even level and media was further mixed with the formaldehyde solution by gentle swirling, followed by a 5 min incubation in the cold room, during which the initial temperature of the media/formaldehyde mixture, being close to room temperature, gradually decreased. To inactivate remaining formaldehyde and the reactive derivatives of fixation 600 µl of a 2.5 M Glycine solution (dissolved in H_2_O), equally chilled, was added to the media/formaldehyde mixture, the same way as described before, followed by another 5 min incubation. Liquid was then aspirated and cells were washed with 25 ml ice cold PBS. The PBS was carefully aspirated and 500 µl of buffer A (10 mM HEPES pH 7.5, 62.5 mM KCl_2_, 2.5 mM MgCl_2_) supplemented with 1 mM DTT, 1 % (w/v) Triton-X-100 and 1x protease inhibitor cocktail (cOmplete mini EDTA-free) was used for in-dish lysis of each two dishes. The lysis buffer was added to the first dish, cells were scraped off quickly and transferred into the 2^nd^ dish together with the lysis buffer. Cells of the 2^nd^ dish were also scraped into the lysis buffer which was then transferred into a 1.5 ml microcentrifuge tube, resulting in a total volume of around 1.8 ml. Cell lysis was allowed to proceed for 10 min more on ice with occasional vortexing. The lysate was clarified by a 5 min centrifugation at 14,000 rpm at 4 ⁰C. The supernatant typically had a concentration of around 10 AU OD_260_ per ml. Aliquots of 10 AU (1 ml) were flash frozen and stored at −80 ⁰C. **Yeast —** Formaldehyde cross-linking was performed according to (Valášek et al., 2007). Culture flasks (no more than two at a time) were put into ice and 250 g of ice were added to the culture followed by swirling to achieve a quick cool down. 27 ml of a 37 % w/v formaldehyde solution were added and promptly mixed with the cells and media, resulting in a total concentration of around 0.8 % w/v, taking the ice volume into account. After 1h of incubation on ice, 50 ml of a 2.5 M glycine solution was added and mixed in. All subsequent steps were performed on ice or 4⁰C. Cells were pelleted by centrifugation, washed with ice-cold buffer B (20 mM Tris/HCl pH 7.4, 200 mM KCl, 5 mM MgAc) and pelleted again. The resulting cell pellet was weighted and resuspended in 1 ml ice-cold complemented buffer B (1 mM DTT, 10 mM PMSF, 1 ug/ml Pepstatin, 1 ug/ml Aprotinin, 1 ug/ml Leupeptin, 1x protease inhibitor cocktail (cOmplete EDTA-free) per 1 g of cells. Cell lysis was performed with a bead beater. A settled bead volume of 500 µl ice-cold acid washed glass beads together with 1 ml cell suspension were jiggled for 40 seconds at speed 5. The supernatant was clarified twice by centrifugation at 14000 rpm for 5 and 10 min. The resulting cell lysate would have a concentration of around 150 U (OD_260_) per ml. Aliquots of 50 U (330 µl) were flash frozen and stored at −80 ⁰C.

#### Footprint generation and 40S/80S separation

**HEK293T—** 1 ml of cell lysate (10 U OD_260_) was incubated with 70 U of RNase I (Invitrogen, cat.no. AM2294) at 24 ⁰C for 30 min with gentle shaking. 28 U of SUPERaseIn (Invitrogen, cat.no. AM2694) were then added and the mixture was loaded onto a 5-45 % (w/v) linear sucrose density gradient (12 ml, in buffer A supplemented with 1 mM DTT). After a high velocity spin (39,000 rpm, 2.5 h, 4 ⁰C, SW41 Ti rotor Beckman-Coulter) the gradients were fractionated and fractions collected. Fractions corresponding to the small ribosomal subunit or the full ribosome, as determined by the UV absorbance traces, were pooled separately..**Yeast—** 330 µl of cell lysate (50 U OD_260_) was incubated with 750 U of RNase I (Invitrogen, cat.no. AM2294) at 24 ⁰C for 1 h with gentle shaking. 100 U of SUPERaseIn (Invitrogen, cat.no. AM2694) were added and 2 aliquots (100 U OD_260_) were loaded onto a 5-30 % (w/v) linear sucrose density gradient (38 ml, in buffer B supplemented with 1 mM DTT). After a high velocity spin (32,000 rpm, 3 h 44 min, 4 ⁰C, SW32 Ti rotor Beckman-Coulter) fractions were collected. Fractions corresponding to the small ribosomal subunit (40S) or the full ribosome (80S) were determined by the UV absorbance profiles and pooled separately.

#### Co-immunoprecipitation (CoIP)

Collected pools of 40S fractions or 80S fractions were divided into a 95 % aliquot, used for the CoIP, and a 5% aliquot that was flash-frozen and used later for TCP-seq library preparation from unselected pools of 40Ses or 80Ses. **HEK293T—** 150 U OD_260_ of cell lysate (corresponding to 15 ‘regular’ aliquots) were used per sample. After RNase I digestion, all aliquots were pooled and loaded onto 12 gradients for the SW41 Ti rotor. After separation, each pool (40S and 80S) was adjusted to 0.07 % (w/v) Triton-X-100 with a 20 % stock solution and split into two. To one half, 100 µl of a 50 % slurry of anti-eIF3b coated protein A/G agarose was added. To the other half, beads without antibody coating were added, to control for the background. Tubes were rotated overnight at 4 ⁰C. Then, beads were collected by spinning at 500 rpm for 3 min at 4 ⁰C and the supernatant was removed. Beads were washed 3 times with 5 ml buffer A supplemented with 0.07 % (w/v) Triton-X-100, and once with buffer A lacking detergent. All supernatant was carefully removed after the last washing step and 600 µl of buffer A were added, before proceeding with TCP-seq library preparation. **Yeast—** To control for the CoIP background, strain SY183 was used, which does not express FLAG-tagged protein. 300 U OD_260_ of cell lysate (corresponding to 6 ‘regular’ aliquots) were used per sample. After RNase I digestion, all aliquots were pooled and loaded onto 3 gradients for the SW28 Ti rotor. After separation, each pool (40S and 80S) was adjusted to 0.2 % Triton-X-100 (w/v) with a 20 % stock solution. 150 µl of a 50 % slurry of ANTI-FLAG M2 affinity gel (Sigma, cat.no. A2220) in buffer B was added and tubes were rotated over night at 4 ⁰C. Beads were spun at 500 rpm for 3 min at 4 ⁰C and supernatant was removed. Beads were washed 4 times with 5 ml buffer B supplemented with 0.2 % (w/v) Triton-X-100 and once with buffer B lacking detergent. Supernatant was carefully removed after the last washing step and elution was carried out by adding 200 µl of a 150 ng/µl FLAG peptide solution, incubating the resultant mixtures for 30 min at 4 ⁰C with gently shaking to keep the beads swirling. Supernatant was used to prepare TCP-seq libraries.

#### Polysome profile analysis

Fifteen A260 units of whole cell extract were separated by high-velocity sedimentation through a 5 % to 45 % sucrose gradient at 39,000 rpm for 2.5 h using the SW41 Ti rotor. The absorbance at 254 nm throughout the gradient was recorded to visualise the ribosomal species. These profiles were recorded on paper, scanned, and further edited to remove the grid and differently colour the traces for better visualisation.

#### TCP-seq library preparation

After the CoIP, RNA was extracted with hot acid phenol method, generally as described by (Ingolia, 2010), from the eluates (yeast) or directly from the beads suspended in 600 ul of buffer A (HEK293T) and the unselected pools of 40S and 80S. Briefly, samples were brought to room temperature and then incubated at 65 ⁰C for 5 min. SDS was added up to a final concentration of 1 % (w/v) using a stock solution of 20 %. One volume of acid phenol-chloroform mix (7:1, v/v) was added, followed by a 20 min incubation at 65 ⁰C with frequent intermittent vortexing. Samples were then placed on ice for 5 min, followed by a 10 min spin at 14,000 rpm in a benchtop Eppendorf centrifuge at 4 ⁰C. A second acid phenol-chloroform (7:1, v/v) extraction was performed at room temperature for 5 min with frequent vortexing. Finally, an extraction with chloroform-isoamylalcohol mix (24:1, v/v) was performed at room temperature and constant vortexing for 1 min. The aqueous phase was recovered by centrifugation and supplemented with 1 µl glycogen (5 mg/ml). Nucleic acids were then precipitated by adjusting to 300 mM sodium acetate (using a 3 M stock solution, pH 5.2) and adding 1 volume of isopropanol. After incubation at −20 ⁰C overnight, the RNA was pelleted by centrifugation, washed with ice-cold 75 % ethanol and air-dried. Pellets were dissolved in 10 µl 10 mM Tris/HCl (pH 8.0). Sequencing libraries were prepared according to (Ingolia et al., 2012), with some adjustments. Importantly, we allowed for a broader size selection which included RNA fragments from around 18 nt up to around 85 nt, except for yeast replicates 2 and 3 used for 80S Sel-TCP-seq (see Table 1) where the size selection was restricted to around 20 −50 nt. Further, for yeast a custom set of rRNA depletion oligos was used according to predominant rRNA fragments found in test libraries.

#### mRNA-seq library preparation

First, total RNA was extracted with the RNeasy Mini Kit. The Poly(A)Purist MAG Kit was further used to extract polyadenylated RNA which following was fragmented with RNA fragmentation reagent for 8 min at 70 ⁰C, wherefore 1ug of RNA was mixed with 1.1 ul of the RNA fragmentation reagent to obtain a final volume of 11.1 ul. The fragmentation was terminated by adding the stop buffer. Subsequent library preparation was performed as for TCP-seq samples according to Ingolia et al. 2012 Nat Protoc, with some adjustments.

#### High-throughput sequencing

All libraries of a replicate and species were multiplexed and sequenced together (see Table 1 for an overview of the used libraries). The human replicate 1 and the yeast replicates 1-1, 1-2 and 1-3 were sequenced with a single end layout and 100 bp read length on the illumina platform HiSeq 2500. The yeast replicates 2 and 3 were sequenced with a single end layout and 50 bp read length on the illumina platform HiSeq 2500. The human replicate 2 was sequenced with a paired end layout and 100 bp read length on the illumina platform HiSeq Xten, but only the first read of the pair was used.

#### RNA sequencing read mapping

First, the adapter sequences were removed from reads, and reads without adapter, adapter-only reads, reads shorter than 18nt after adapter removal and reads of low quality were discarded, using fastx_clipper and fastq_quality_filter from the FASTX-Toolkit. Reads were further filtered by successive alignment to ribosomal RNA (using bowtie), a variety of ncRNAs and tRNA (using Bowtie2), retaining unaligned reads for further analyses. tRNA sequences were retrieved from the tRNA database of the University Leipzig (http://trna.bioinf.uni-leipzig.de). Using the default settings of Tophat, the resulting filtered pool of reads was first aligned to the appropriate transcriptome; reads that did not align, were mapped to the appropriate genome. For ***S. cerevisiae***, the genome assembly R64-1-1 was used and a custom annotation file was created containing all entries with the gene_biotype protein_coding of the Ensemble annotation R64-1-1.80 and further 5’ and 3’ UTR annotation as well as 5’ UTR intron annotation retrieved from the SGD database in November 2017 (whereby the longest annotated UTRs per gene were added to the transcriptome). For genes without annotated 5’ or 3’ UTRs, 50nt of the adjacent genomic sequence were added as generic UTRs. For ***H. sapiens***, the genome assembly GRCh38 and the Ensembl annotation GRCh38.80 were used.

#### tRNA enrichment analysis

Primary alignments of reads that mapped to the tRNA reference were used. Counts for all tRNA isoforms decoding for the same amino acid were combined with the distinction of initiator and elongator methionine tRNAs. The ratio of counts derived from FOI::40S libraries versus counts from unselected 40S libraries were plotted in Figures 3J,K and S10D.

#### Metagene heatmaps

Primary read alignments were used. Only start or stop codons were used which are the sole annotated start or stop codon in all annotated transcripts tagged ‘protein_coding’ of a gene in the Ensembl annotations GRCh38.80 and R64-1-1.80 for human and yeast, respectively, resulting in 6639 start sites and 6514 stop sites for yeast. For human, further, only start sites were considered whose corresponding genes have a minimal annotated 5’UTR and CDS length of 80 nt and 30 nt, respectively, which corresponds to the depicted 5’UTR and CDS range. Likewise, only stop sites were considered whose corresponding genes have a minimal annotated CDS and 3’UTR length of 80 nt and 30 nt, respectively. To increase the number of used start codons, in case a gene has more than one start site annotated, the mainly used start site was included if it could be reliably appointed from the unselected 40S libraries (A start codon got appointed the mainly used one if its FP coverage comprised 75% of the sum of the coverage of all annotated start codons of a gene or it comprised 60% of all but was at least 3 times higher than the second highest. The FP coverage in the area of −3 to +5 nucleotides from the A of the start codon was used (with A=0).). This resulted in altogether 10778 start sites and 16803 stop sites for human.

#### Assigning reads to a transcript feature

Primary alignments were used. To reliably determine the feature of a transcript only genes were considered with a sole start and stop codon annotated for all transcripts tagged ‘protein_coding’ of that gene in the Ensembl annotations GRCh38.80 and R64-1-1.80 for human and yeast, respectively, yielding 9241 genes for human and 5818 genes for yeast (whereby for yeast also genes tagged ‘dubious open reading frame’ and ‘unlikely to encode a functional protein’ in the SGD gene description (August 2018) were excluded). In general, 10 nt were added upstream and downstream the 5’UTR and 3’UTR, respectively, owing to our observation that FPs often extend some nucleotides beyond the annotated transcript ends. If no UTRs were annotated for a gene, 50 nt for yeast and 100 nt for human upstream the start codon and/or downstream the stop codon were used as 5’ and 3’UTRs, respectively. Assigning FPs to transcript areas: 5’UTR: The FP 3’ end should map upstream of +5 nt of the A of the start codon and the FP 5’ end should map on or downstream the transcript start. Start codon: The 5’end should map on or upstream of −3 nt of the A of the start codon and the 3’ end should map on or downstream of +5. CDS: The 5’ end should map downstream of −3 nt of the A of the start codon and the 3’ end should map upstream of −6 of the first nucleotide of the stop codon. Stop codon: The 5’ end should map on or upstream of −6 nt of the first nucleotide of the stop codon and the 3’ end should map on or downstream of +2. 3’UTR: The FP 5’ end should map downstream of +2 of the first nucleotide of the stop codon. Always, both ends should map to an exon. Violin plots in Figures 3G-I, 3L, S10C and S10E were plotted with default settings of the function violinplot() from the Python library seaborn, Python Software Foundation. Python Language Reference, version 2.7. Available at http://www.python.org).

#### Footprint 3’ end distributions in relation to the start codon

Primary alignments were used. Start sites were considered being the sole annotated start site of all transcripts tagged ‘protein_coding’ of a gene if this gene has a minimal annotated 5’ UTR of 50 nt (11174 start sites for human and 5994 start sites for yeast). FPs whose 5’ end map within −14 to −12 of a start site (with A = 0) were included for figures 2D,E and S8E-G. FPs assigned to a start codon (see ‘*Assigning reads to a transcript feature*’) were considered for figure 4A. 1000 iterations of random resampling were performed (see QUANTIFICATION AND STATISTICAL ANALYSES for details).

#### Count per gene for correlation plots

Unique alignments were used. Count was performed with htseq-count (mode ‘union’, feature type ‘exon’ and id attribute ‘gene_id’). For human the Ensembl annotation GRCh.38.80 was used. For yeast Ensemble annotation R64-1-1.80 was used including further the 5’ and 3’ UTR annotation as well as the 5’ UTR intron annotation from the SGD database in November 2017 (whereby the longest annotated UTRs per gene were added to the transcriptome). For genes without annotated 5’ or 3’ UTRs, 50nt of the adjacent genomic sequence were added as generic UTRs. Generally, only genes tagged ‘protein_coding’ were included.

#### Confirming transcript starts for metagene plots with cap alignment

mRNA datasets were used. Transcritps with a minimal total count of 100 FPs should have an average coverage of maximal 3 in the region from −30 to −10 of the annotated transcript start and an average coverage of minimal 20 in the region from +10 to +30 of the annotated transcript start to be considered a confirmed transcript start. Further, the annotated 5’ UTR should have a minimal length of 80 nt to avoid interference of the start codon associated FPs with long 5’ extensions. This yielded 190 and 136 transcript starts for yeast and human, respectively.

#### Transcriptome-wide screen for co-translational assembly in the 80S Sel-TCP-seq data

The average coverage of two intervals within the CDS of a gene was compared. Interval 1 was located in the beginning of the CDS from +30 up to +90 nt (with A of the start codon set = 0). Interval 2 was located in the end of the CDS from −90 up to −30 nt (with the first nucleotide of the stop codon set = 0). A gene was identified as candidate for co-translational assembly, if the average coverage of interval 2 was at least 5 times higher than that of interval 1. Nine candidates were found. Five of those were dropped because further investigation revealed a FLAG like epitope in the protein sequence of the gene in the right distance to the onset of the increase in coverage (*RPA34, RTF1, RNH203, CDC16, LEU1*). Coverage tracks of unselected and FOI bound 80S FPs for the other four candidates were analysed further and are shown in Figure 7.

#### Calculation of the fold induction to estimate the co-translational binding

The average footprint coverage after the onset of induction to the end of the CDS was divided by the average coverage before the onset starting from the beginning of the CDS. The first and last 30 nt of the CDS were not regarded as well as 30 nt +/- the onset position. The fold induction seen in the FOI::80S Sel-TCP-seq data was normalized by the fold induction seen in the unselected 80S TCP-seq data. The normalization was done to account for features in the mRNA which might slow down or stall translating ribosomes, leading to a coverage increase not attributable to co-translational binding.

### QUANTIFICATION AND STATISTICAL ANALYSES

Statistical details of experiments can be found in the figure legends and figures. One and Two sample t-tests were performed using the R function t-test. A p value larger than 0.05 is referred to as not significant (n.s.). Random gene wise resampling was performed for data shown in figures 2D, 2E, 4A and S8E-G to estimate the influence of individual genes on the density profiles. This was done by randomly re-selecting genes, allowing duplications, *n* times (where *n* is the original sample size i.e. the number of genes used, as indicated in each figure legend). The mean and 95^th^ percentiles of such iterations are shown in the figures. A custom Python script was used.

### DATA AND CODE AVAILABILITY

The datasets generated during this study were deposited in the GEO database with the following accession number: (The number will be inserted once the data was deposited.) All custom Python scripts used for the analyses in this paper are available upon request.

### KEY RESOURCES TABLE

**SUPPLEMENTAL ITEMS**

**Table S1. Yeast strains used in this study.**

