## supplemental figures for "Selective Translation Complex Profiling Reveals Staged Initiation and Co-translational Assembly of Initiation Factor Complexes"

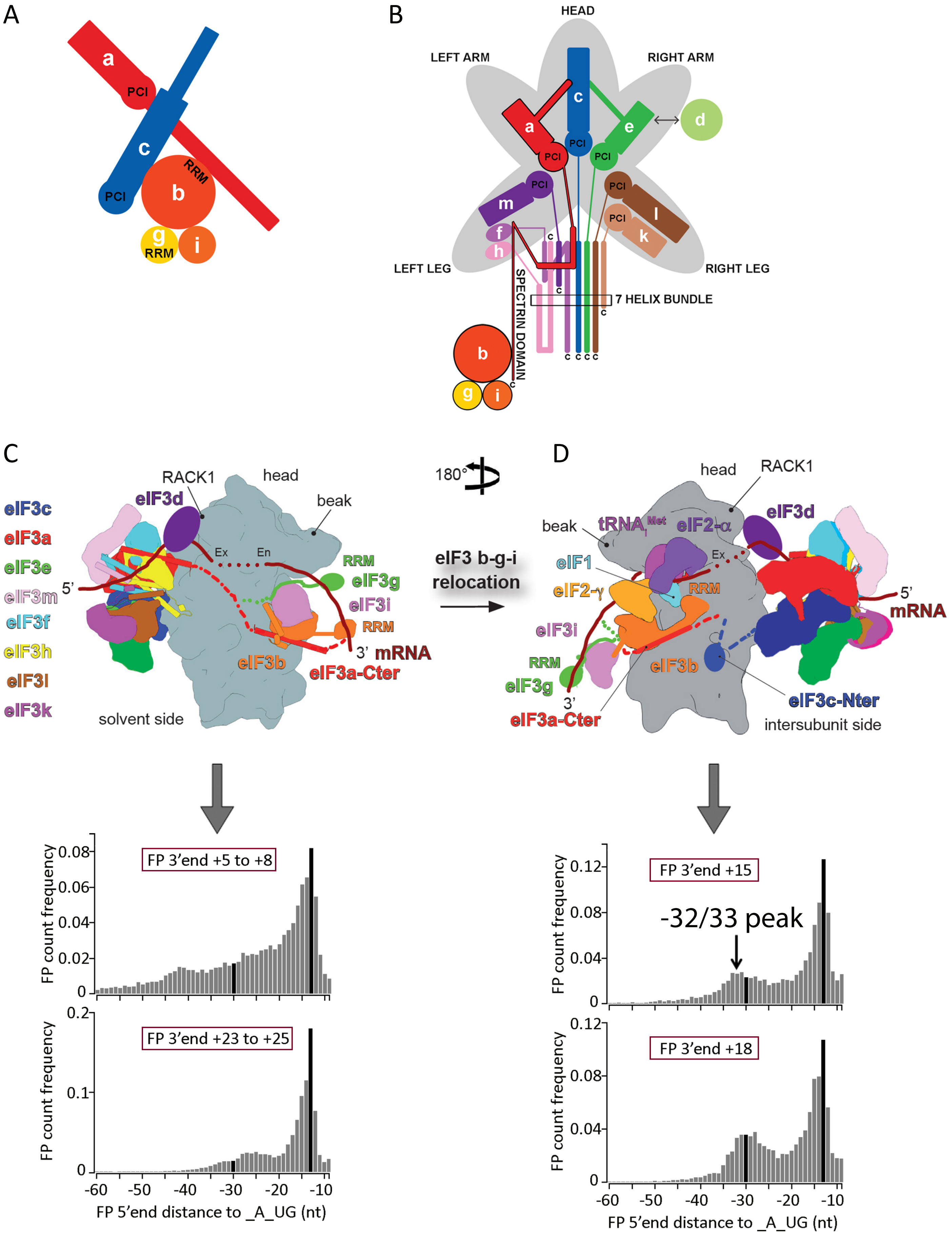
A
a
PCI
c
RRM
b
PCI
g
RRM
i
B
HEAD
LEFT ARM
RIGHT ARM
c
a
e
d
PCI
PCI
PCI
PCI
m
PCI
l
PCI
k
f
h
LEFT LEG
RIGHT LEG
7 HELIX BUNDLE
SPECTRIN DOMAIN
b
g
i
C
RACK1
head
beak
eIF3d
eIF3c
eIF3a
eIF3e
eIF3m
eIF3f
eIF3h
eIF3l
eIF3k
5'
Ex
En
RRM
eIF3g
eIF3i
RRM
3' mRNA
eIF3b
eIF3a-Cter
solvent side
180°
eIF3 b-g-i relocation
D
head
RACK1
beak
tRNAi Met
eIF2-α
eIF3d
eIF1
Ex
eIF2-γ
RRM
5'
mRNA
eIF3i
RRM
eIF3g
3'
eIF3b
eIF3a-Cter
eIF3c-Nter
intersubunit side
FP count frequency
FP 3'end +5 to +8
FP 3'end +23 to +25
FP 3'end +15
-32/33 peak
FP 3'end +18
0.08
0.06
0.04
0.02
0
0.2
0.1
0.12
-60
-50
-40
-30
-20
-10
FP 5'end distance to _A_UG (nt)

A

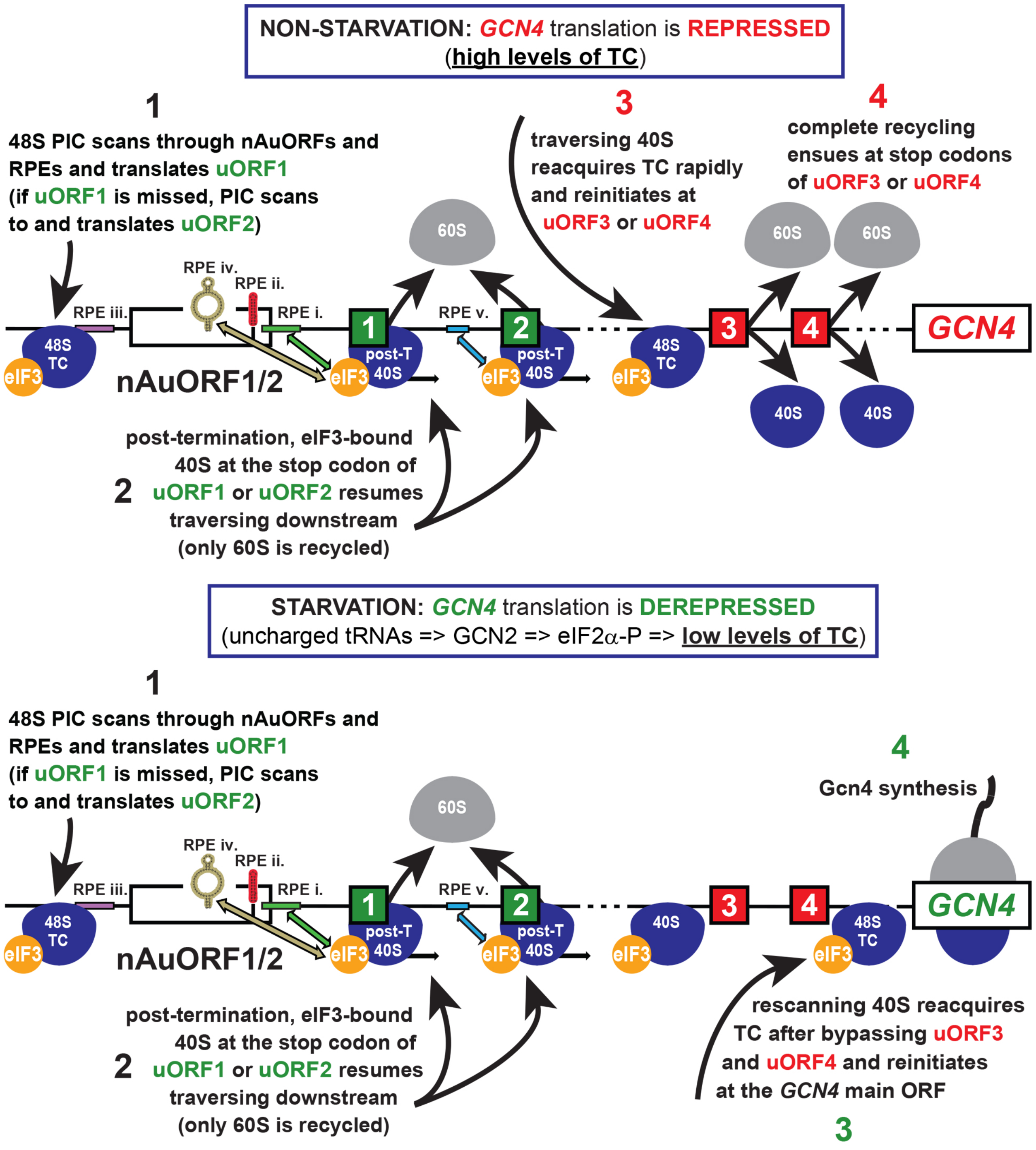

NON-STARVATION: GCN4 translation is REPRESSED
(high levels of TC)
1
48S PIC scans through nAuORFs and RPEs and translates uORF1 (if uORF1 is missed, PIC scans to and translates uORF2)
3
traversing 40S reacquires TC rapidly and reinitiates at uORF3 or uORF4
4
complete recycling ensues at stop codons of uORF3 or uORF4
60S
RPE iv.
RPE ii.
RPE iii.
RPE i.
RPE v.
48S TC
eIF3
nAuORF1/2
post-T 40S
40S
GCN4
post-termination, eIF3-bound 40S at the stop codon of
2 uORF1 or uORF2 resumes traversing downstream
(only 60S is recycled)
STARVATION: GCN4 translation is DEREPRESSED
(uncharged tRNAs => GCN2 => eIF2α-P => low levels of TC)
4
Gcn4 synthesis
rescanning 40S reacquires TC after bypassing uORF3 and uORF4 and reinitiates at the GCN4 main ORF
3


B

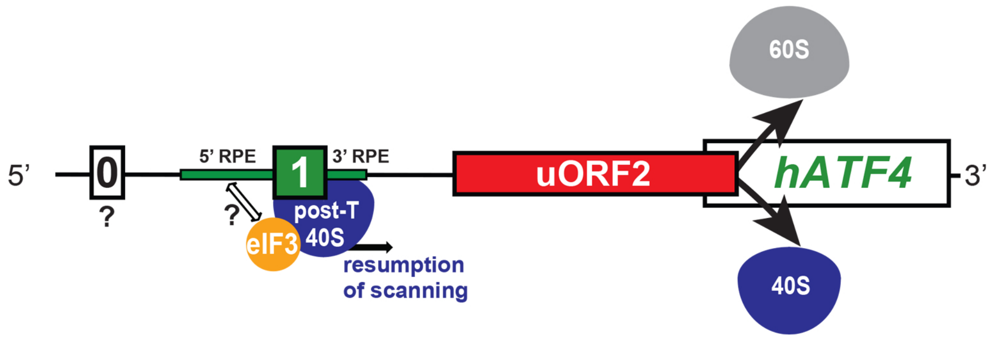

60S
5'
0
?
5' RPE
1
3' RPE
post-T 40S
eIF3
resumption of scanning
uORF2
hATF4
3'
40S

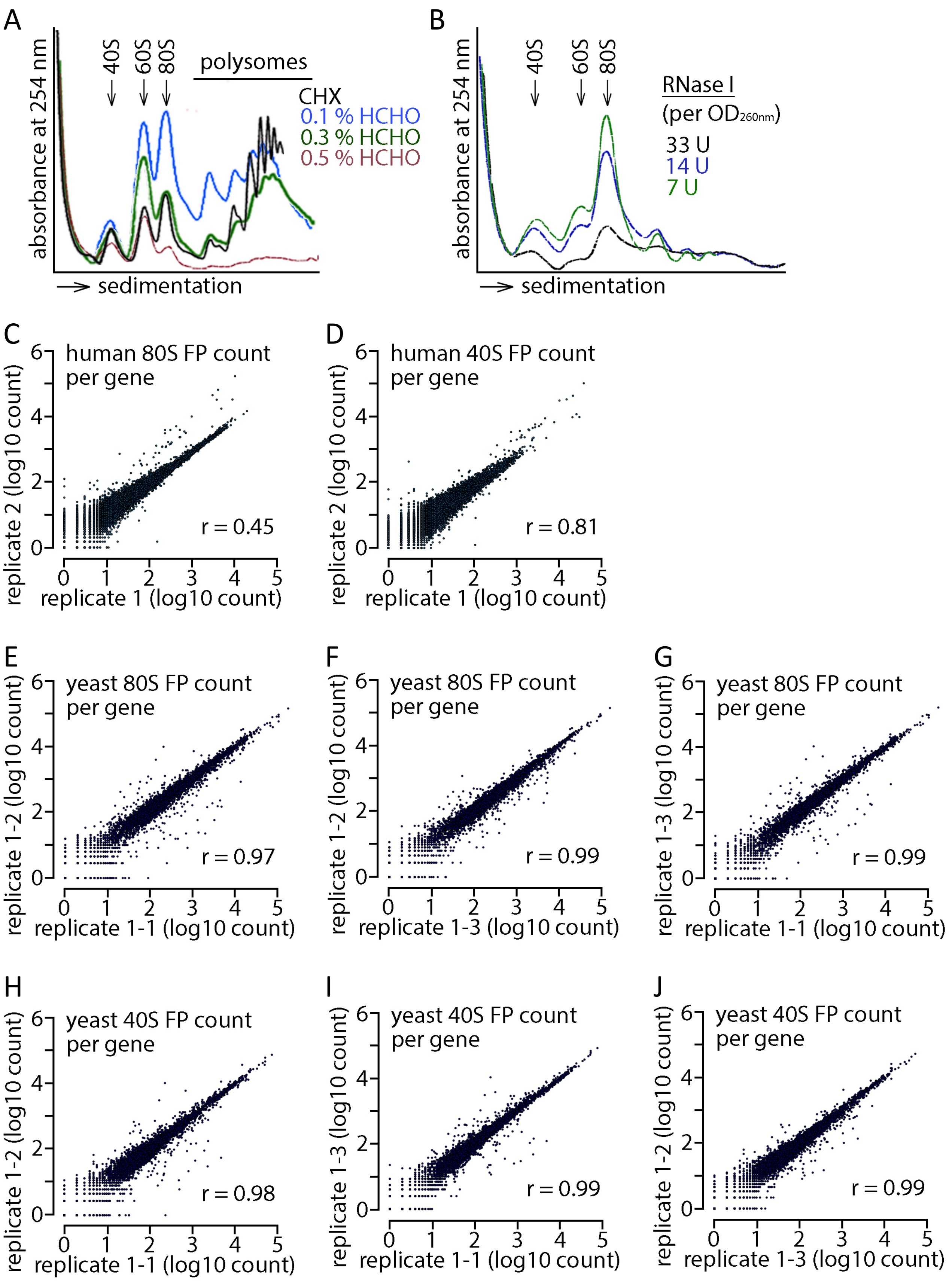
A
absorbance at 254 nm
40S
60S
80S
polysomes
CHX
0.1 % HCHO
0.3 % HCHO
0.5 % HCHO
sedimentation
B
absorbance at 254 nm
40S
60S
80S
RNase I
(per OD260nm)
33 U
14 U
7 U
sedimentation
C
human 80S FP count per gene
replicate 2 (log10 count)
replicate 1 (log10 count)
r = 0.45
D
human 40S FP count per gene
replicate 2 (log10 count)
replicate 1 (log10 count)
r = 0.81
E
yeast 80S FP count per gene
replicate 1-2 (log10 count)
replicate 1-1 (log10 count)
r = 0.97
F
yeast 80S FP count per gene
replicate 1-2 (log10 count)
replicate 1-3 (log10 count)
r = 0.99
G
yeast 80S FP count per gene
replicate 1-3 (log10 count)
replicate 1-1 (log10 count)
r = 0.99
H
yeast 40S FP count per gene
replicate 1-2 (log10 count)
replicate 1-1 (log10 count)
r = 0.98
I
yeast 40S FP count per gene
replicate 1-3 (log10 count)
replicate 1-1 (log10 count)
r = 0.99
J
yeast 40S FP count per gene
replicate 1-2 (log10 count)
replicate 1-3 (log10 count)
r = 0.99

Wagner_Figure S4

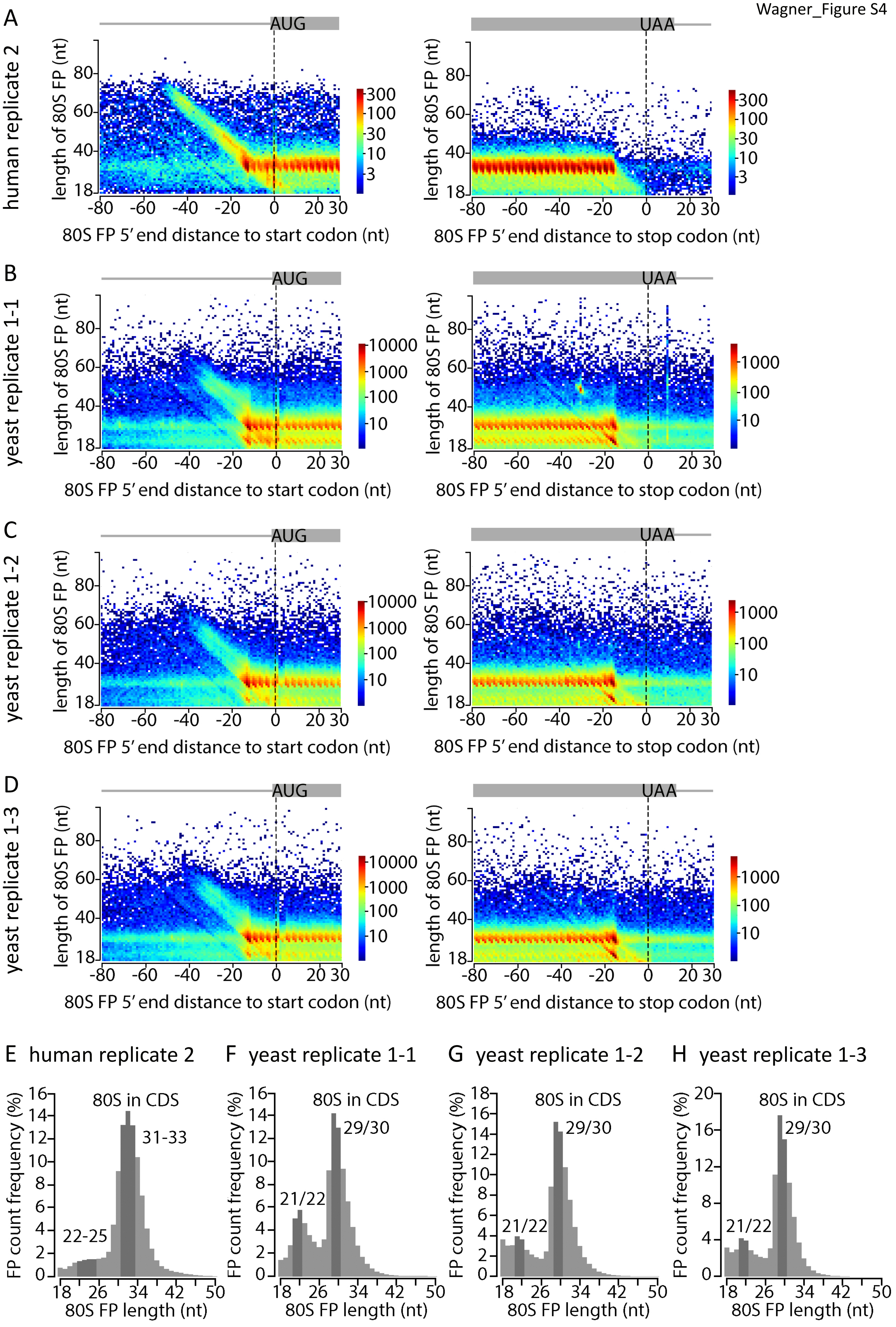

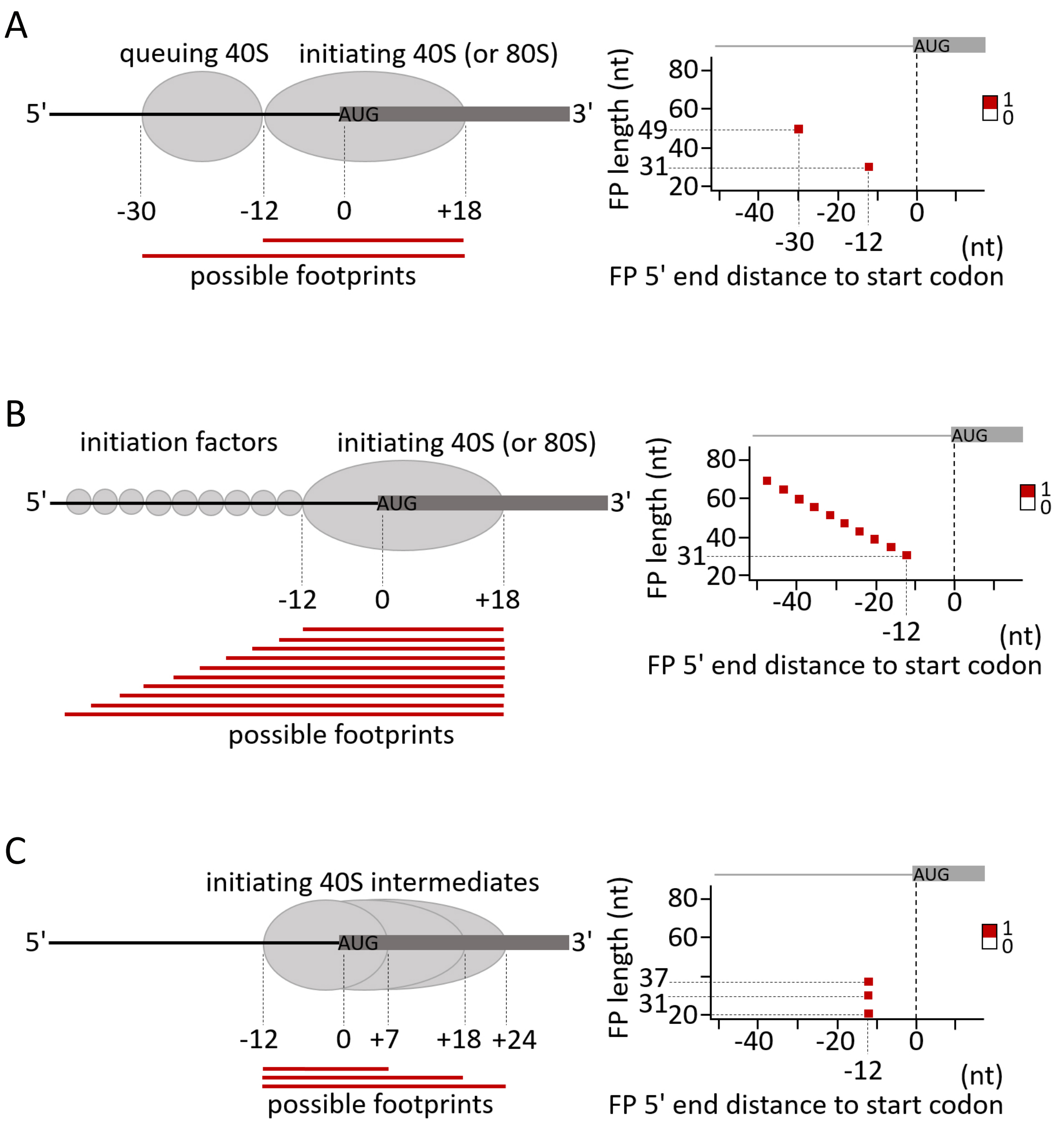
A
queuing 40S
initiating 40S (or 80S)
5'
AUG
3'
-30
-12
0
+18
possible footprints
FP length (nt)
80
60
49
40
31
20
-40
-20
0
-30
-12
(nt)
FP 5' end distance to start codon
1
0
B
initiation factors
initiating 40S (or 80S)
5'
AUG
3'
-12
0
+18
possible footprints
FP length (nt)
80
60
40
31
20
-40
-20
0
-12
(nt)
FP 5' end distance to start codon
1
0
C
initiating 40S intermediates
5'
AUG
3'
-12
0
+7
+18
+24
possible footprints
FP length (nt)
80
60
37
31
20
-40
-20
0
-12
(nt)
FP 5' end distance to start codon
1
0

Wagner_Figure S6

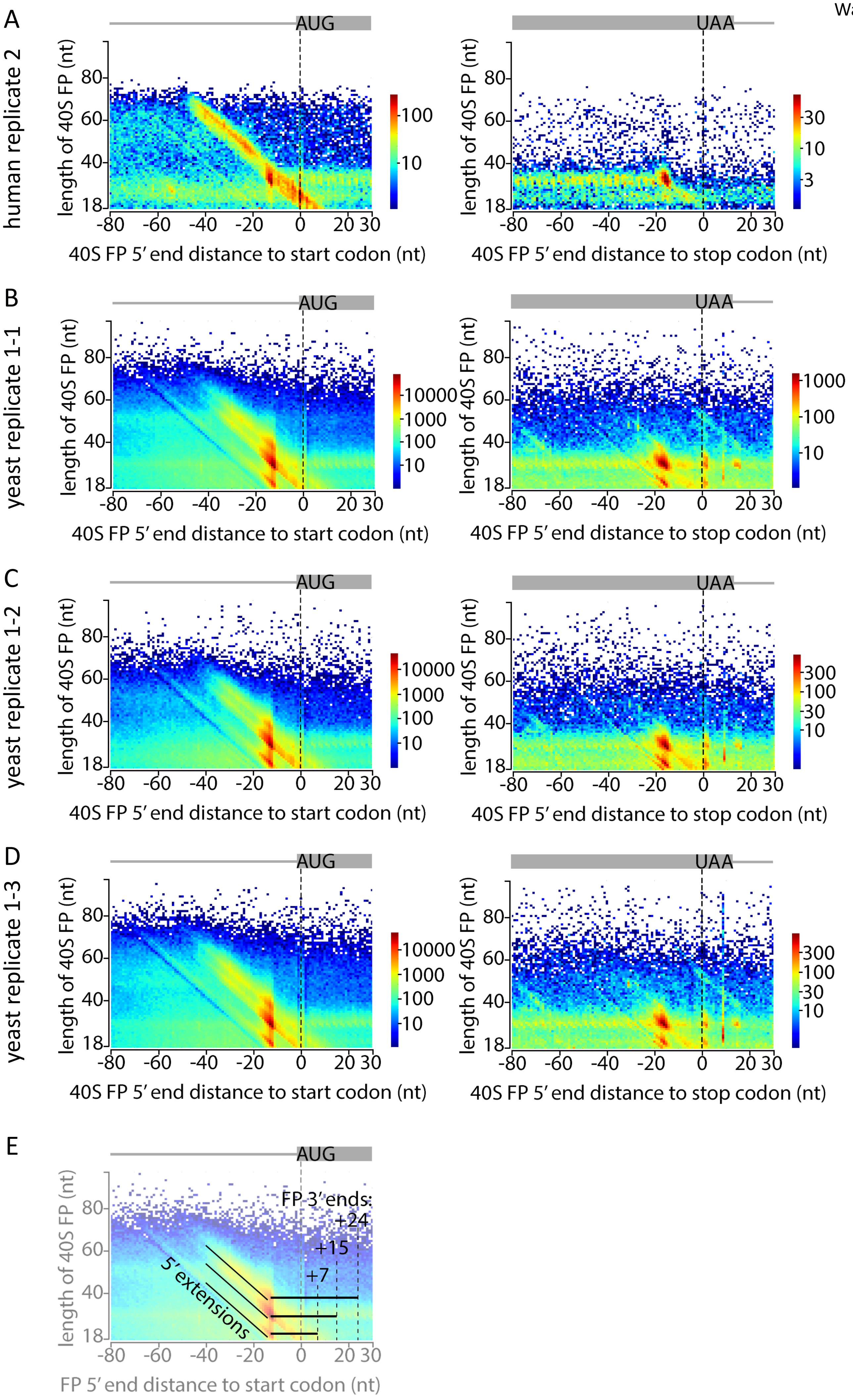

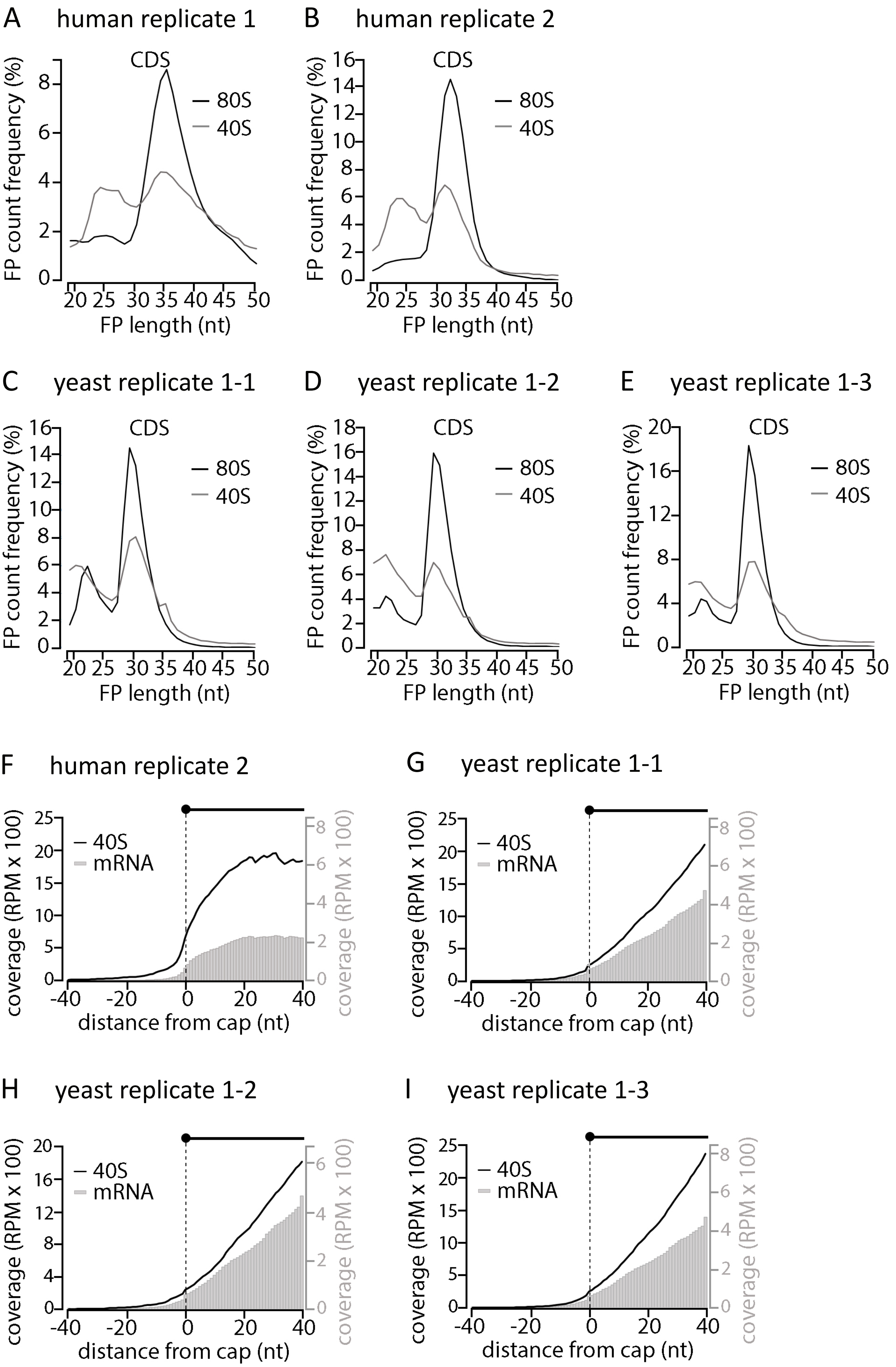
A human replicate 1
CDS
80S
40S
FP count frequency (%)
FP length (nt)
B human replicate 2
C yeast replicate 1-1
D yeast replicate 1-2
E yeast replicate 1-3
F human replicate 2
40S
mRNA
coverage (RPM x 100)
distance from cap (nt)
G yeast replicate 1-1
H yeast replicate 1-2
I yeast replicate 1-3

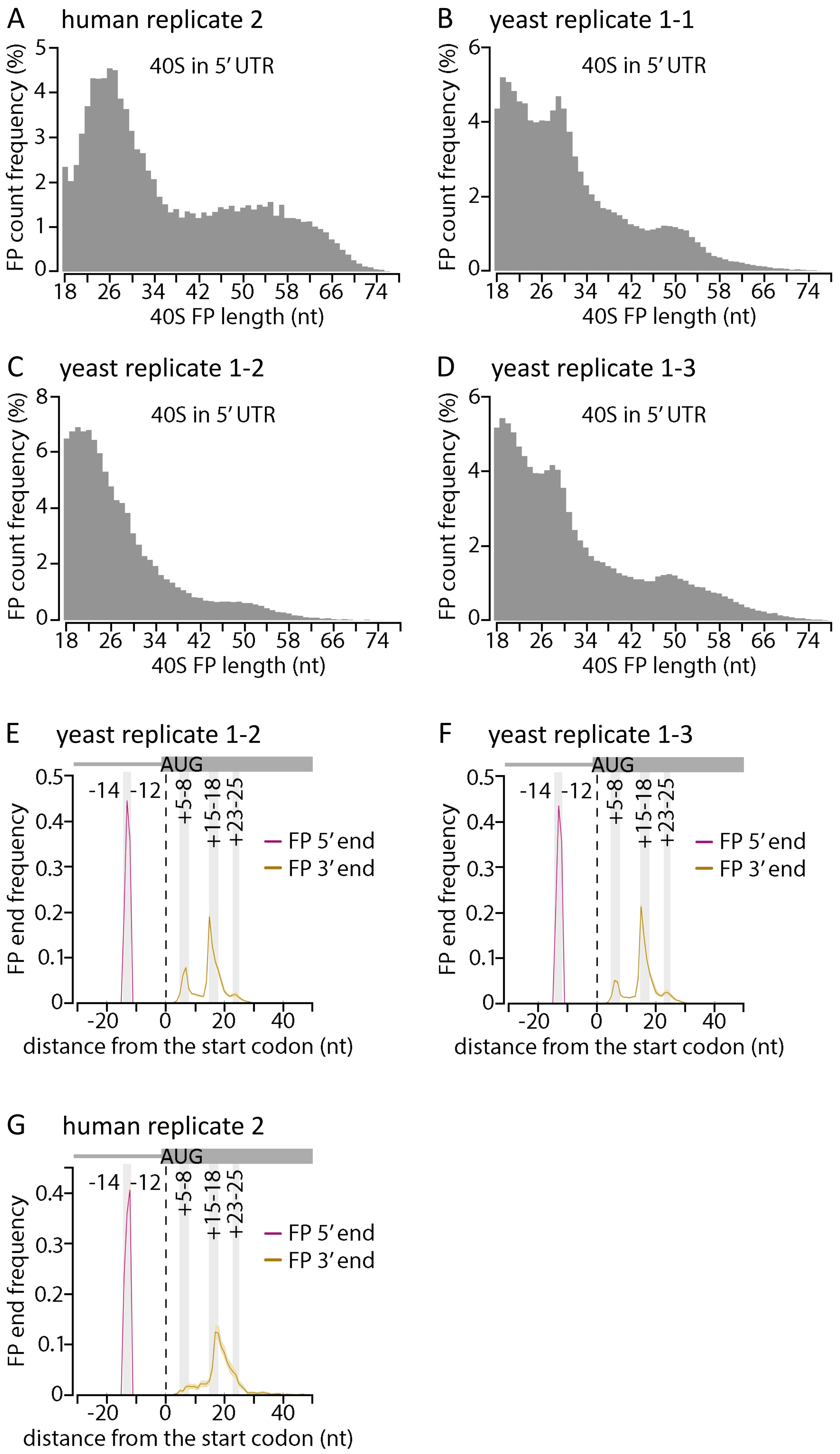
A
human replicate 2
40S in 5′ UTR
FP count frequency (%)
40S FP length (nt)
B
yeast replicate 1-1
40S in 5′ UTR
FP count frequency (%)
40S FP length (nt)
C
yeast replicate 1-2
40S in 5′ UTR
FP count frequency (%)
40S FP length (nt)
D
yeast replicate 1-3
40S in 5′ UTR
FP count frequency (%)
40S FP length (nt)
E
yeast replicate 1-2
AUG
-14 -12
+5-8
+15-18
+23-25
FP 5′ end
FP 3′ end
FP end frequency
distance from the start codon (nt)
F
yeast replicate 1-3
AUG
-14 -12
+5-8
+15-18
+23-25
FP 5′ end
FP 3′ end
FP end frequency
distance from the start codon (nt)
G
human replicate 2
AUG
-14 -12
+5-8
+15-18
+23-25
FP 5′ end
FP 3′ end
FP end frequency
distance from the start codon (nt)

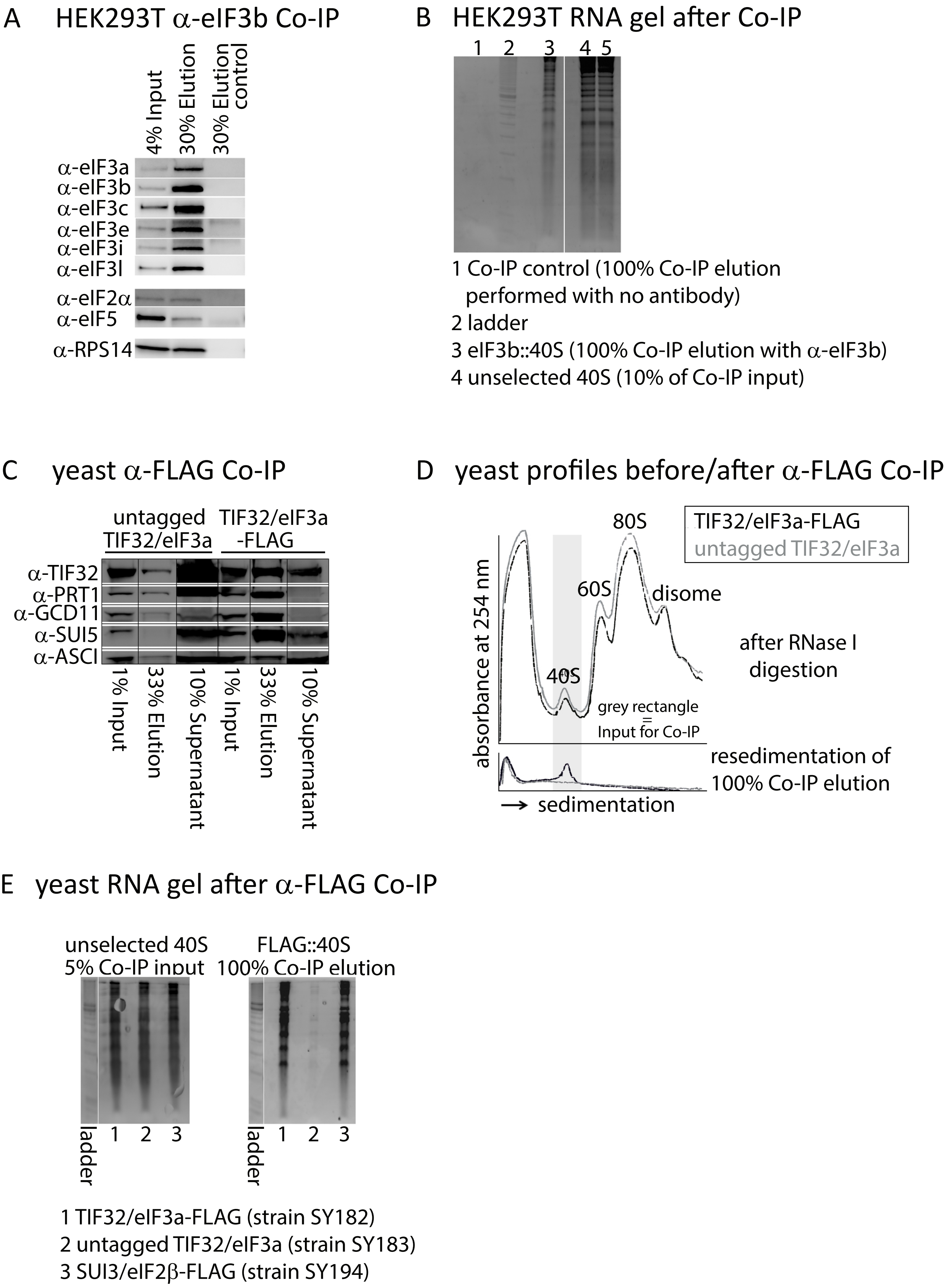
A HEK293T α-eIF3b Co-IP
4% Input
30% Elution
30% Elution control
α-eIF3a
α-eIF3b
α-eIF3c
α-eIF3e
α-eIF3i
α-eIF3l
α-eIF2α
α-eIF5
α-RPS14
B HEK293T RNA gel after Co-IP
1 2 3 4 5
1 Co-IP control (100% Co-IP elution performed with no antibody)
2 ladder
3 eIF3b::40S (100% Co-IP elution with α-eIF3b)
4 unselected 40S (10% of Co-IP input)
C yeast α-FLAG Co-IP
untagged TIF32/eIF3a
TIF32/eIF3a -FLAG
α-TIF32
α-PRT1
α-GCD11
α-SUI5
α-ASCI
1% Input
33% Elution
10% Supernatant
1% Input
33% Elution
10% Supernatant
D yeast profiles before/after α-FLAG Co-IP
TIF32/eIF3a-FLAG
untagged TIF32/eIF3a
80S
60S
disome
40S
after RNase I digestion
grey rectangle = Input for Co-IP
absorbance at 254 nm
resedimentation of 100% Co-IP elution
→ sedimentation
E yeast RNA gel after α-FLAG Co-IP
unselected 40S 5% Co-IP input
FLAG::40S 100% Co-IP elution
ladder 1 2 3
ladder 1 2 3
1 TIF32/eIF3a-FLAG (strain SY182)
2 untagged TIF32/eIF3a (strain SY183)
3 SUI3/eIF2β-FLAG (strain SY194)

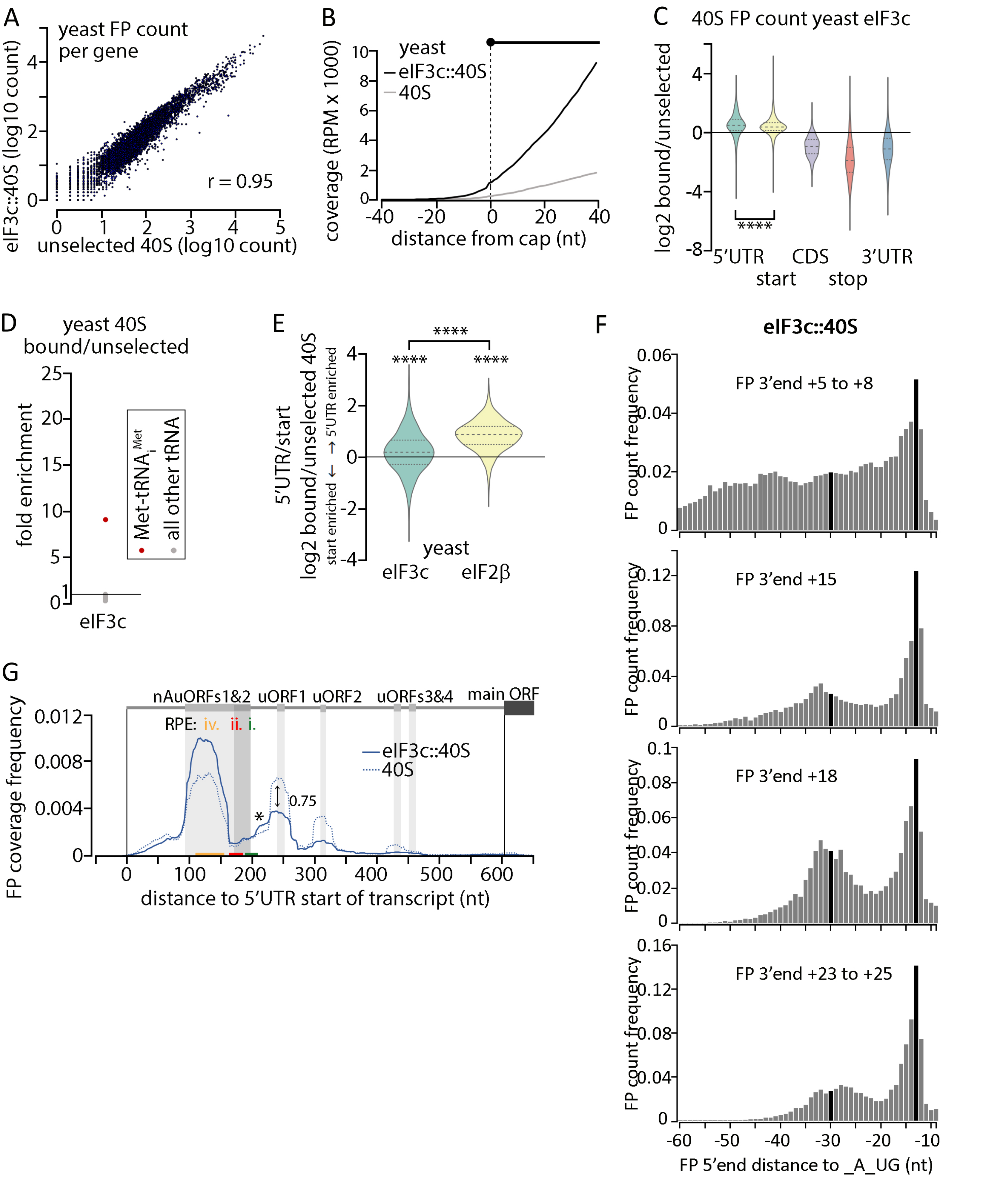
A
yeast FP count per gene
eIF3c::40S (log10 count)
unselected 40S (log10 count)
r = 0.95
B
yeast
eIF3c::40S
40S
coverage (RPM x 1000)
distance from cap (nt)
C
40S FP count yeast eIF3c
log2 bound/unselected
****
5'UTR
start
CDS
stop
3'UTR
D
yeast 40S bound/unselected
fold enrichment
Met-tRNAi Met
all other tRNA
eIF3c
E
5'UTR/start
log2 bound/unselected 40S
start enriched ← → 5'UTR enriched
****
yeast
eIF3c
eIF2β
F
eIF3c::40S
FP 3'end +5 to +8
FP 3'end +15
FP 3'end +18
FP 3'end +23 to +25
FP count frequency
FP 5'end distance to _A_UG (nt)
G
nAuORFs1&2
uORF1
uORF2
uORFs3&4
main ORF
RPE: iv. ii. i.
eIF3c::40S
40S
0.75
*
FP coverage frequency
distance to 5'UTR start of transcript (nt)

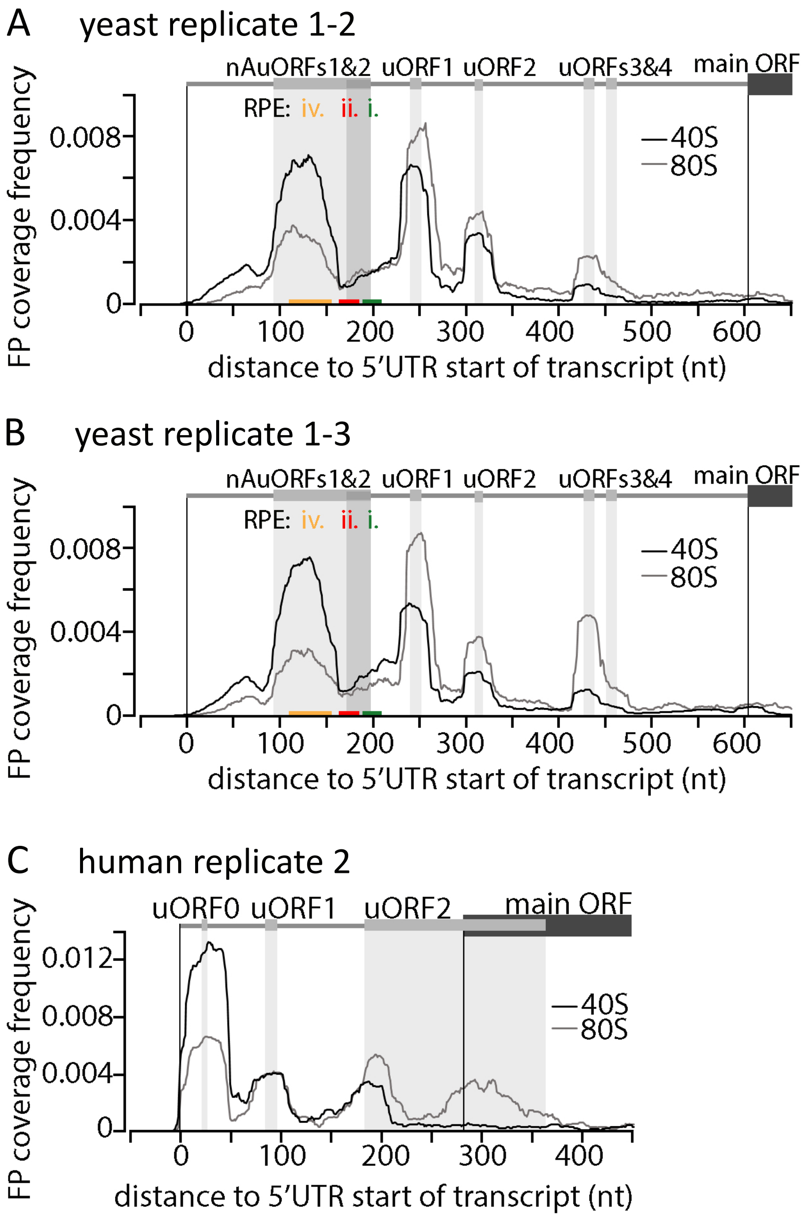
A
yeast replicate 1-2
nAuORFs1&2 uORF1 uORF2 uORFs3&4 main ORF
RPE: iv. ii. i.
40S
80S
FP coverage frequency
0.008
0.004
0
0 100 200 300 400 500 600
distance to 5'UTR start of transcript (nt)
B
yeast replicate 1-3
nAuORFs1&2 uORF1 uORF2 uORFs3&4 main ORF
RPE: iv. ii. i.
40S
80S
FP coverage frequency
0.008
0.004
0
0 100 200 300 400 500 600
distance to 5'UTR start of transcript (nt)
C
human replicate 2
uORF0 uORF1 uORF2 main ORF
40S
80S
FP coverage frequency
0.012
0.008
0.004
0
0 100 200 300 400
distance to 5'UTR start of transcript (nt)

A

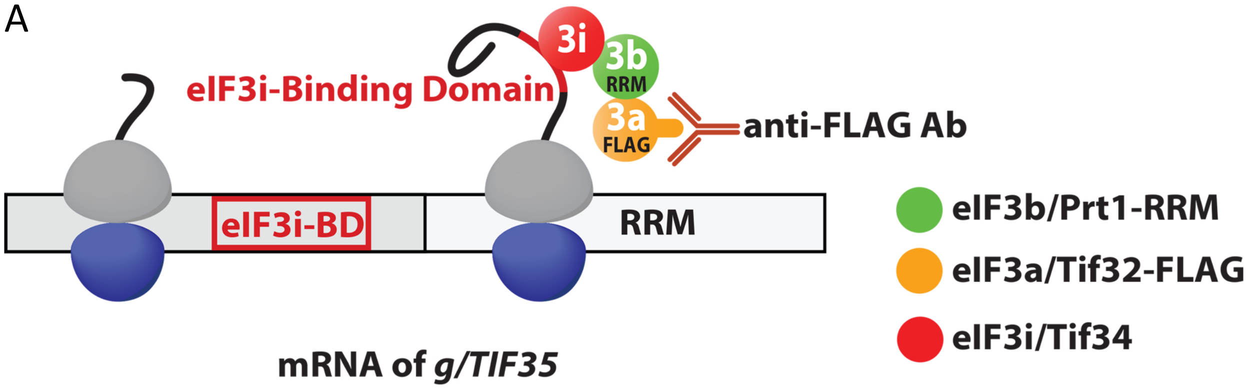
eIF3i-Binding Domain
3i
3b
RRM
3a
FLAG
anti-FLAG Ab
eIF3i-BD
RRM
mRNA of g/TIF35
eIF3b/Prt1-RRM
eIF3a/Tif32-FLAG
eIF3i/Tif34

B

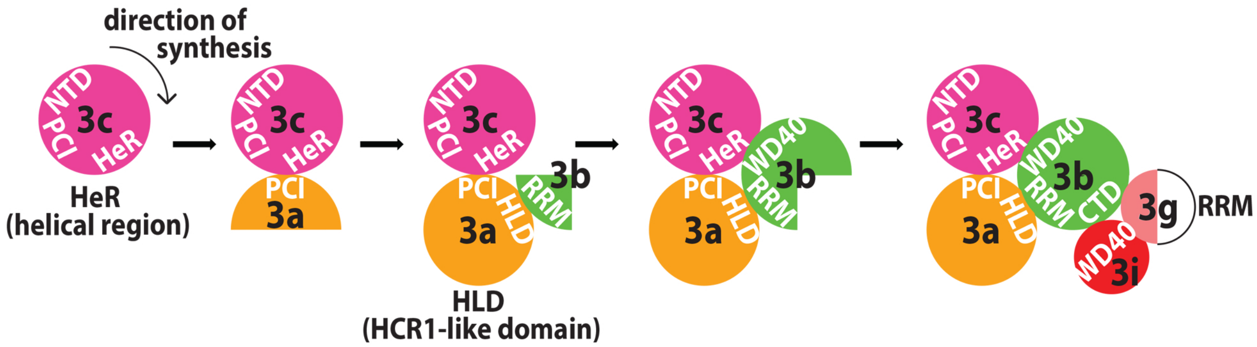
direction of synthesis
NTD
PCI
3c
HeR
HeR
(helical region)
3a
3b
RRM
HLD
HLD
(HCR1-like domain)
WD40
CTD
3g
RRM
3i

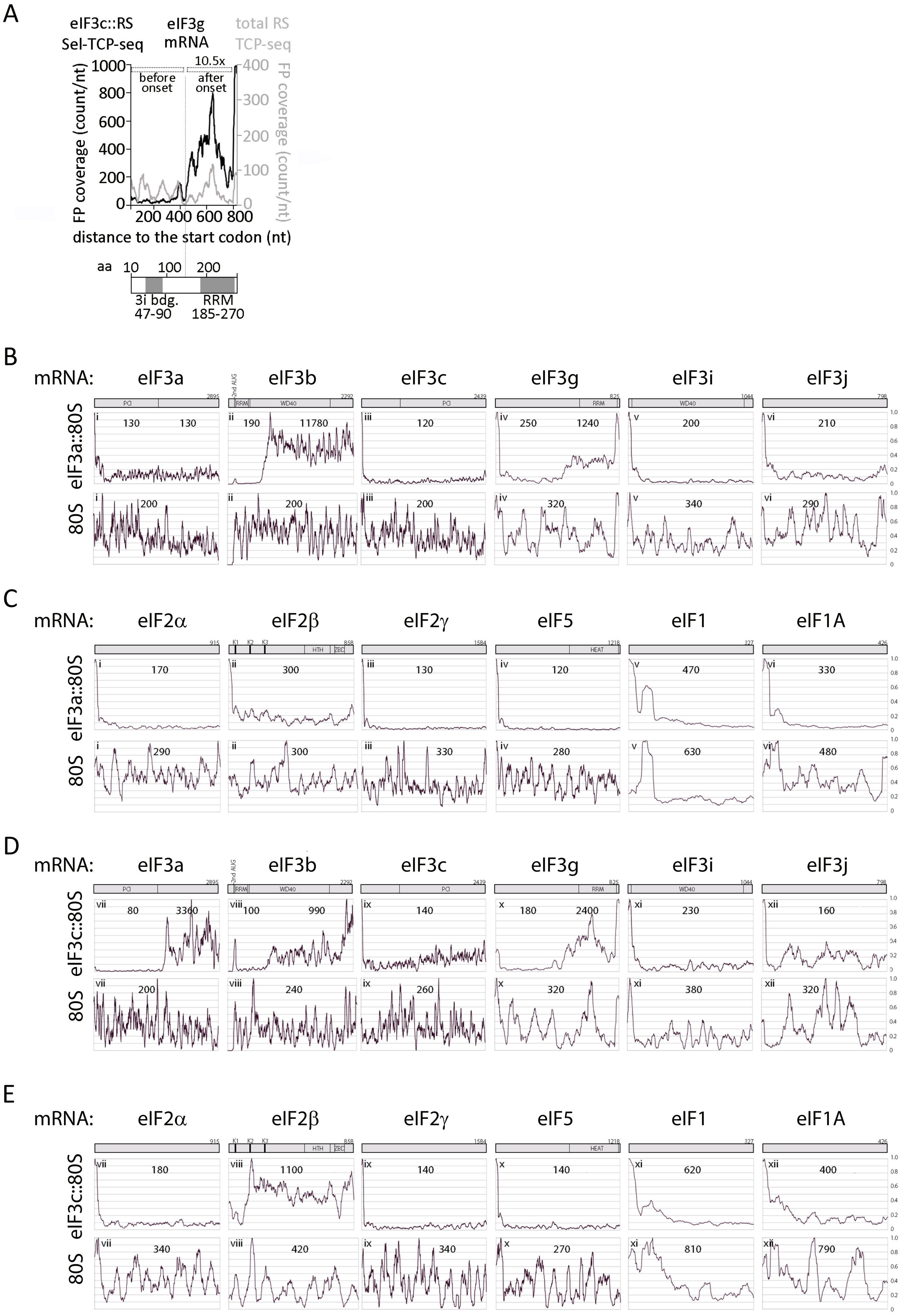
A
eIF3c::RS Sel-TCP-seq
eIF3g mRNA
total RS TCP-seq
FP coverage (count/nt)
before onset
after onset
10.5x
distance to the start codon (nt)
aa 10 100 200
3i bdg. 47-90
RRM 185-270
B
mRNA: eIF3a eIF3b eIF3c eIF3g eIF3i eIF3j
eIF3a::80S
80S
C
mRNA: eIF2α eIF2β eIF2γ eIF5 eIF1 eIF1A
eIF3a::80S
80S
D
mRNA: eIF3a eIF3b eIF3c eIF3g eIF3i eIF3j
eIF3c::80S
80S
E
mRNA: eIF2α eIF2β eIF2γ eIF5 eIF1 eIF1A
eIF3c::80S
80S
